# Mechanosensory Systems and Sensory Integration Mediate *C. elegans* Negative Gravitaxis

**DOI:** 10.1101/2022.03.03.482913

**Authors:** Caroline Ackley, Lindsey Washiashi, Neda Ziaei Kajbaf, Ruchira Krishnamurthy, Zhenxuan Sun, Vivian Duong, Kaylin Choe, Elijah Lane, Cricket Wood, Eleanor Smith, Giulia Pellegrini, Mark Sherwin, Pradeep Joshi, Joel H. Rothman

## Abstract

The ability to sense Earth’s gravitational pull is essential for orientation, navigation, and proprioception in many organisms. We report here that *C. elegans* exhibits pronounced negative gravitaxis, or movement away from the Earth’s center, independent of orientation of the geomagnetic field. This behavior is antagonized by light and electromagnetic fields, suggesting that it is integrated with other sensory inputs. We found that MEC-5/7/12, and TRPA-1, but not MEC-4/10 DEG/ENaC channels or other proteins involved in gentle touch transduction, are essential for negative gravitaxis, suggesting that the sensory system for detecting and responding to gravity is separable from touch sensation. We also found that the PVD neurons but not touch receptor neurons (TRNs) are required for this behavior. These findings implicate an interconnected mechanism for gravity sensation involving an ion channel that is also present in the mammalian vestibular system, suggesting possible homology in gravity sensing across animal phylogeny.

## Introduction

Gravity sensation is a common trait among most Eukaryotes. Members of the protists, fungi, plants, and animals depend on gravity sensation for survival. Small changes in the position or orientation of these organisms result in a mechanical force that is transduced by graviperceptive organelles or organs. This force is often conveyed through dense organelles or mineral-rich structures whose displacement triggers a signaling pathway that ultimately results in a behavioral output ***Ross et al. (1984***). A common characteristic of gravity transduction pathways in animals is the use of ciliated neurons, in which deflection of hair-like “stereocilia” opens mechanically gated ion channels ***Bezares-Calderón et al. (2020***); ***Lacquaniti et al. (2014***). Although this general mechanism has been well characterized, less is known about how mechanoreceptors and mechanosensory cells transduce such minute forces – estimated to be as small as 0.57-1.13 piconewtons (pN) in single-celled Euglena, for example (which lack stereocilia) – into a robust signal ***Häder and Hemmersbach (2017***).

Given the availability of a detailed connectome of the entire nervous system and powerful genetic and optogenetic tools, *C. elegans* has been a highly effective model organism for elucidating the neural circuitry and molecular mechanisms governing sensory perception, learning, memory, and behavior. However, because they are traditionally studied in a two-dimensional environment on the surface of Petri dishes containing agar, perpendicular to the vector of gravity, few studies have examined complex behavior exhibited by *C. elegans* in three dimensions ***Guisnet et al. (2021***). In the wild, textitC. elegans is typically found in moist compost, such as under shrubs or along riverbanks, which can change drastically from season to season ***Frézal and Félix (2015***); ***Kodama-Namba et al. (2013***). These animals have evolved an adaptive alternative larval phase, the dauer larvae, that prioritizes dispersal over reproduction during adverse environmental conditions. Dauer larvae exhibit characteristic nictation, in which they “stand” on their tails and wave their heads. This behavior is believed to facilitate dispersal, possibly by allowing worms to “hitch a ride” on larger animals that pass by, such as isopods ***Lee et al. (2012***). Therefore, gravitational force may be a critical input that *C. elegans* use in combination with other cues to navigate to the surface before traveling longer distances ***Figure 1***.

**Figure 1.**
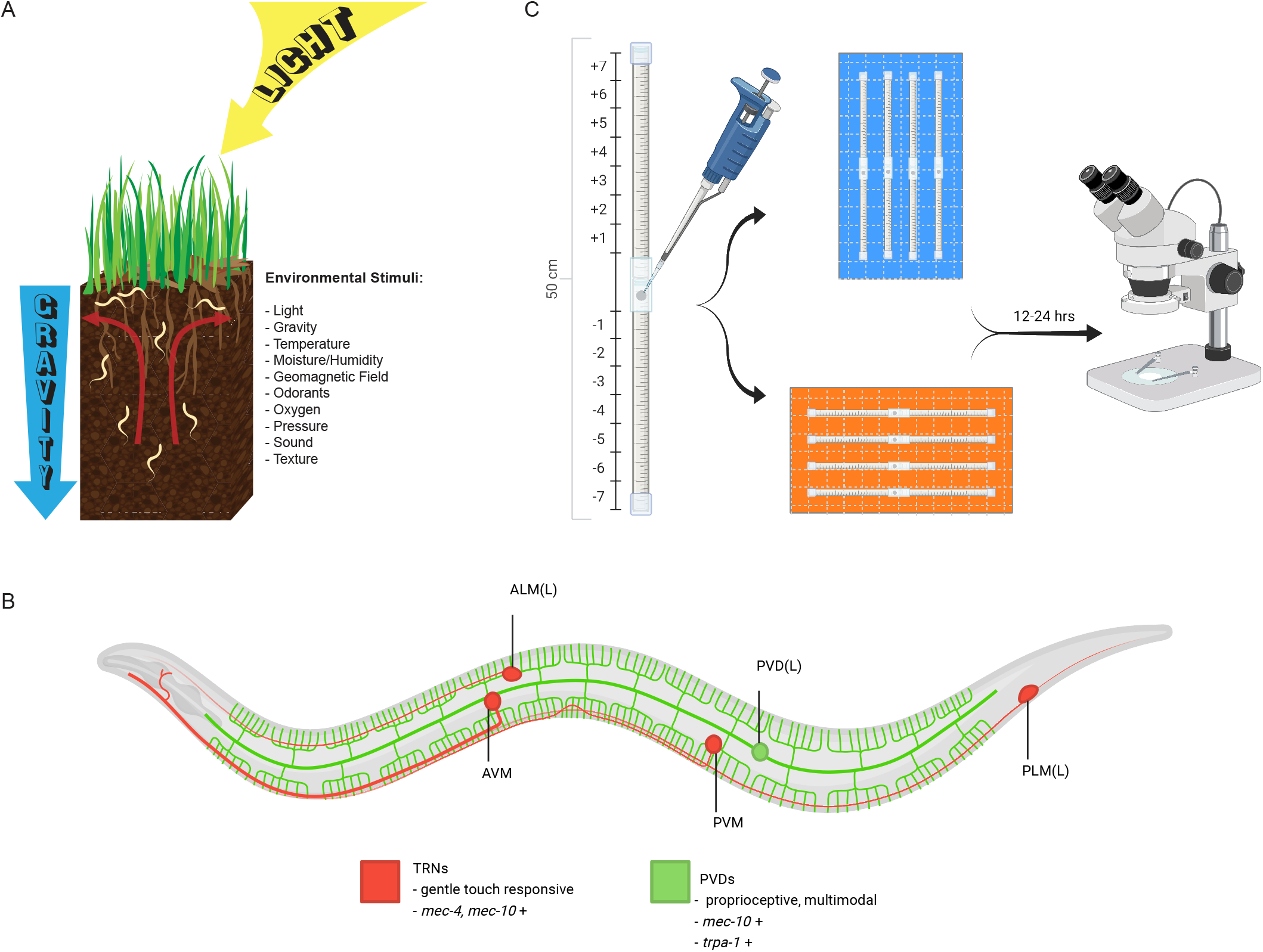
Behavioral model, wiring diagram, and gravitaxis assay design. **(A)** Graphic depicting environmental stimuli that inform *C. elegans* behavior in a natural setting. **(B)** Diagram of key neurons mentioned in this study. TRNs are labeled in red and PVD neurons in green (for neurons with left and right pairs, only the left neuron is displayed). Adapted from individual neuron illustrations available on WormAtlas ***Altun (n.d***). **(C)** Gravitaxis chamber and experimental design used in this study. Figures **(B)** and **(C)** created with BioRender.com.

In a previous study, it was reported that *C. elegans* dauer larvae showed no gravitactic preference, in contrast with *C. japonica*, which negatively gravitaxes on vertically oriented Petri plates ***Okumura et al. (2013***). Other studies reported that *C. elegans* show a tendency to orient downwards when swimming in liquid, suggesting potential positive gravitaxis ***Chen et al. (2021***). However, as in *Drosophila* and the ascidian *Ciona*, the ability of nematodes to undergo gravitaxis is likely context-dependent ***Bae et al. (2016***); ***Bostwick et al. (2020***). *C. elegans* shows a strong aversive response to light ***Ward et al. (2008***) and are highly responsive to electrical fields ***Gabel et al. (2007***); ***Rezai et al. (2010***); ***Sukul and Croll (1978***). Magnetotaxis has also been observed in these worms ***Vidal-Gadea et al. (2015***), although the reproducibility of this behavior has been challenged ***Landler et al. (2018b***,a); ***Malkemper et al. (2023***); ***Njus et al. (2015***); ***Vidal-Gadea et al. (2018***). Integration with these or other sensory modalities may influence or mask gravitactic behavior and experiments to test for gravitaxis in the absence of light or electromagnetic fields (EMF) have no been reported with *C. elegans*. In this study we report that in the absence of light, *C*.*elegans* dauers exhibit pronounced negative gravitaxis, which is strongly enhanced when they are shielded from ambient electromagnetic fields and light.

While little is known about their behavioral responses to gravity, the effects of hyper- and microgravity on *C. elegans* physiology has been documented ***Gao et al. (2015***, 2017); ***Kalichamy et al. (2016***); ***Saldanha et al. (2016***); ***Xu et al. (2014***). In response to hypergravitational forces, signaling by the mechanosensory DEG/ENaC sodium channel proteins, MEC-4/MEC-10, leads to nuclear localization of the DAF-16 FoxO transcription factor, which also transduces insulin-like growth factor signaling and stress responses ***Kim et al. (2007***). The *mec-4/10* genes in *C. elegans*, which encode channel proteins, and several other “mec” genes involved in gentle touch sensation, are expressed in the six touch receptive neurons (TRNs), which specifically mediate gentle touch sensation ***Chalfie and Sulston (1981***). While the MEC-4 and -10 proteins function together in the TRNs, they also show non-overlapping expression in some cells, such as the proprioceptive PVD neurons ***Figure 1B***. Mechanosensation in many animals is also mediated by TRP (transient receptor potential) cation channels, which are common across metazoan phylogeny ***Chatzigeorgiou et al. (2010***); ***Han et al. (2013***); ***Kindt et al. (2007***); ***Montell (2003***). The TRPA-1 receptor is a polymodal sensor, capable of conveying either high or low temperatures as well as light and noxious stimuli ***Venkatachalam et al. (2014***). In worms, TRPA-1 confers sensitivity to noxious cold in PVD neurons as well as mechanosensation in OLQ and IL1 neurons ***Han et al. (2013***). The *trpa-1* homologs *pyx* and *pain* are necessary for gravitaxis in *Drosophila* ***Sun et al. (2009***).

In this study, we sought to determine whether *C. elegans* possesses a system for detecting normal gravitational force and whether known mechanosensory molecular components and neurons participate in response to gravity. We discovered that *C. elegans* prefers to migrate vertically against the force of gravity – i.e., to undergo negative gravitaxis. Further, we found that gravitaxis is profoundly influenced by environmental cues of light and background electromagnetic fields, revealing sensory integration in the behavior. We also report that the MEC-7/12 microtubule components and MEC-5 collagen protein, but not the associated MEC-4/10 DEG/ENaC channel proteins or several other components involved in TRN function and touch sensation, are required for gravitactic behavior. Further, we found that the TRPA-1 channel is essential for negative gravitaxis in *C. elegans*. Finally, we found that the TRPA-1 channel is essential for negative gravitaxis in *C. elegans*. Finally, we identified the polymodal PVD neurons, which span the length of the worm, as essential for this directional bias. Based on these findings, we propose a model of gravity transduction that involves the unique action of MEC-5/7/12, TRPA-1, and PVD neurons. Our findings suggest that even in a diminutive animal with low mass, a potentially homologous system for gravity sensing integrates with other sensory inputs to optimize responses to environmental cues and its orientation on the planet.

## Results

### *C. elegans* preferentially migrates vertically against the force of gravity

We investigated whether *C. elegans* can detect and respond to gravity by initially focusing on the behavior of the dispersal state, the dauer larva. When first stage (L1) larvae experience stressful conditions, including overcrowding, lack of resources, and extreme temperatures, they subsequently develop into an alternative third-stage larva, the dauer larva. Through pronounced physical, metabolic, and behavioral changes, dauers become efficient vectors for dispersal that can survive for months without food ***Wang et al. (2009***). Dauer larvae of many nematodes show nictation behavior, in which they raise their heads at a 90° angle to the surface plane, raising the possibility that they can orient in the gravitational field, though this behavior may simply reflect orientation perpendicular to the surface.

Under normal laboratory growth conditions in which *C. elegans* are cultivated on flat agar surfaces in Petri dishes, dauers of the laboratory N2 strain of *C. elegans* do not nictate unless contaminated by fungi or when grown on three-dimensional habitable scaffolds ***Guisnet et al. (2021***); however, 3-D scaffolds are not convenient for studying migratory behavior. To address this issue, we adapted a setup used to study neuromuscular integrity ***Bainbridge et al. (2016***); ***Beron et al. (2015***) and magnetotaxis ***Vidal-Gadea et al. (2015***) in *C. elegans* to investigate whether *C. elegans* dauers exhibit directional bias in response to gravity ***Ackley et al. (2022***) (see Methods). We injected dauers into the gravitaxis assay chambers comprised of two serological pipettes that, when stacked end-to-end, allow for ∼25 cm of movement in either direction from the injection site ***Figure 1C***. Chambers loaded with worms were oriented vertically to test for movement in the gravitational field and migration of each individual was scored 12-24 hours later. Positive gravitaxis (migration between bins -1 to -7) or negative gravitaxis (+1 to +7) was determined by comparing the distribution against paired normal distributions centered about the origin (see Methods). In several experiments, we assayed horizontally oriented chambers simultaneously to control for any directional preference attributable to the construction and design of the chamber itself.

In contrast to prior published assays conducted on Petri plates, which failed to reveal gravitactic preference with *C. elegans* ***Okumura et al. (2013***), we found that N2 dauer larvae showed a weak but significant directional bias in migration toward the top of the vertical chamber under normal laboratory conditions compared to simulated normal distributions and horizontal controls (average vertical location = +1.28, as defined in Methods, n =2,222 worms over 10 trials; p <0.0001, vertical vs. Normal Distribution^*a*^ and Normal Distribution^*b*^; p <0.001, vertical vs. horizontal; Kruskal-Wallis test; ***Figure 2A–B***). For comparison, we applied a “Gravitaxis index” (analogous to the Chemotaxis Index ***Larsch et al. (2015***); ***Srinivasan et al. (2012***)) using the equation 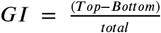 and performed a two-sided t test against a hypothetical mean of 0. The results of this test were marginally significant, demonstrating the difficulty of assessing gravitaxis using a traditional behavioral index (GI vertical = 0.43 +/-0.17, SEM). Together, these observations suggest that *C. elegans* dauer larvae show a mild preference for upward migration under normal laboratory conditions when assayed across a migration field that is >5x larger than that of previous studies ***Okumura et al. (2013***).

**Figure 2.**
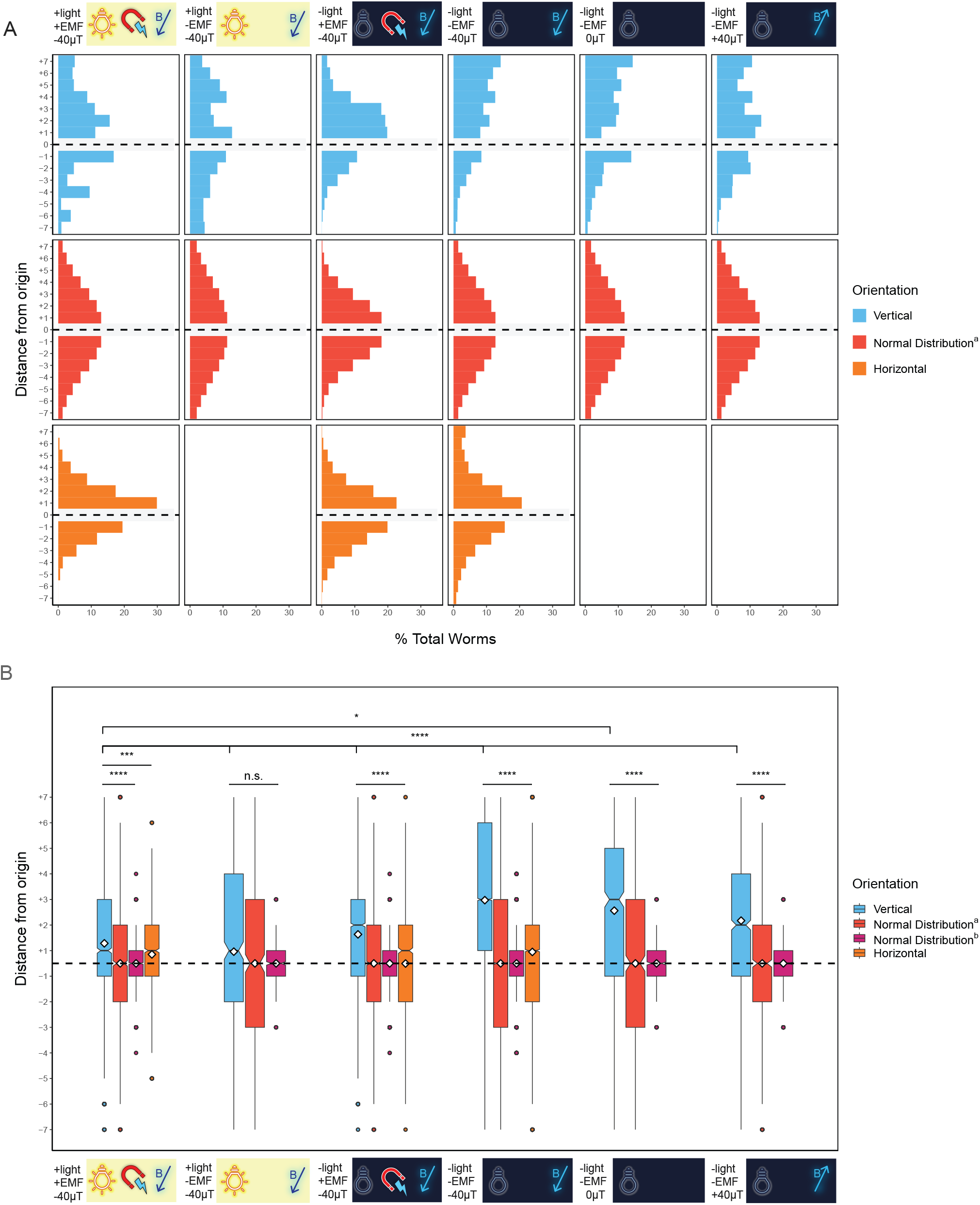
*C. elegans* negatively gravitax under different environmental conditions. **(A)** Histograms depicting cumulative distribution of *C. elegans* worms over multiple trials in vertical (blue) and horizontal (orange) assays as well as in comparison with a simulated normal distribution (Normal Distribution^*a*^), which is created using a mean of 0 and the same standard deviation as the combined vertical assay distributions. Horizontal grey box and dotted line indicate the center of the assay, which was not scored. Presence or absence of normal overhead light and background EMF (no Faraday cage) are denoted with +/-. -40*μ*T indicates normal exposure to Earth’s magnetic field, which is ∼ -40*μ*T in Santa Barbara, CA. **(B)** Boxplots depicting data shown in A. Dauers tested in a dark Faraday cage are used in future comparisons. * p <0.05, ** p <0.01, *** p <0.001, **** p <0.0001; n.s. is not significant using Kruskal-Wallis followed by Dunn’s test with Bonferroni correction. Notches on boxplots represent 95% confidence intervals; mean values are indicated with a diamond. Normal Distribution^*a*^ is described above; Normal Distribution^*b*^ was created using a mean of 0 and the same sample size as the combined vertical assays. Dotted horizontal line indicates the origin of the assay (location “0”), which was not scored. **Figure 2—figure supplement 1**. Faraday cage integrity influences gravitactic behavior **Figure 2—figure supplement 2**. Solenoid coil assay setup **Figure 2—figure supplement 3**. Worms oriented parallel or perpendicular to Earth’s magentic field also gravitax **Figure 2—figure supplement 4**. AB1 dauers isolated in Adelaide, Australia migrate up despite inverted local magnetic field inclination **Figure 2—figure supplement 5**. A negative gravitactic trend is also observed in ***Landler et al. (2018b***) **Figure 2—figure supplement 6**. Gravitaxis behavior occurs independently from thermotaxis along shallow temperature gradients

### Evidence for sensory integration: negative gravitaxis is attenuated by light and alternating electromagnetic fields

External and internal sensory cues that provide important context alter behavioral responses in animals ***Chen and Chalfie (2014***); ***Sengupta (2013***). As. *C. elegans* exhibits negative phototactic behavior ***Ward et al. (2008***), we posited that negative gravitaxis might be adversely influenced by this response to light ***Figure 1A***. To address this possibility, we repeated the gravitaxis assay in the dark using a light-blocking blackout cloth. In sharp contrast to the lack of discernible directional preference in the horizontal chambers, we found that N2 dauer larvae demonstrated an enhanced and highly significant preference for upward migration in the vertical chambers under these conditions (average vertical location = +1.63, n = 4,358 worms, 12 trials; p <0.0001, all comparisons) ***Figure 2A–B***. This preference for upward migration was significantly greater (p <0.0001) than that seen under normal laboratory lighting conditions ***Figure 2B***, suggesting that light attenuates the ability of the animals to sense or respond to the force of gravity.

*C. elegans* has also been reported to respond to direct ***Chrisman et al. (2016***); ***Gabel et al. (2007***); ***Sukul and Croll (1978***) and alternating ***Rezai et al. (2010***) currents, as well as magnetic fields ***Vidal-Gadea et al. (2015***). To test whether EMF, as with light, influences the gravitactic behavior of dauers, we analyzed their movement in chambers that were shielded from the ambient EMF present in the lab using a Faraday cage, which blocks EMF produced by alternating currents but not the magnetic field that is generated by the direct current of the Earth’s dynamo (see Methods). We found that shielding the chambers from both EMF and light resulted in dramatically enhanced negative gravitactic behavior: in such chambers, dauers showed strong negative gravitaxis, with highly significant directional bias compared to horizontal controls and simulated normal distributions (average vertical location = +2.97, n = 34,766 worms, 142 trials; p <0.0001, all comparisons; ***Figure 2A–B***). Further, the animals showed significantly stronger negative gravitaxis under these conditions than either the -light +EMF, +light -EMF, or +light +EMF conditions, when comparing the vertical distributions of all worms assayed (p <0.0001 in all cases). Over the course of several months of use, we noticed that gravitaxis in our Faraday cage gradually decreased as a result of diminished EMF blocking, as confirmed by restoration of cell phone reception in the cage (see Methods). Strong negative gravitatic behavior was restored after reinforcing the cage ***figure Supplement 1***. Thus, the ability of *C. elegans* to undergo negative gravitaxis is altered by both light and EMF, suggesting that gravitaxis behavior is integrated with other sensory inputs by the animal. In all subsequent experiments, we used a Faraday cage in the dark to block these other sensory inputs.

### Vertical movement is not in response to Earth’s magnetic field or temperature gradients

Previous studies have debated the evidence for magnetosensation in *C. elegans* ***Landler et al. (2018a***,b); ***Malkemper et al. (2023***); ***Njus et al. (2015***); ***Vidal-Gadea et al. (2015***, 2018) While an early report showed that worms undergo magnetotaxis ***Vidal-Gadea et al. (2015***), subsequent investigators in several publications have presented data refuting this claim ***Landler et al. (2018a***,b); ***Malkemper et al. (2023***); ***Njus et al. (2015***). Despite this controversy, we sought to determine whether the upward migration of the animals that we observed might be attributable to geomagnetism. As Faraday cages do not effectively shield against Earth’s magnetic field generated by direct current, we built a solenoid coil to mitigate magnetic field strength along the vertical axis (*B*_*z*_) to determine whether magnetotaxis is responsible for upward movement. We assayed taxis behavior in a Faraday cage within the solenoid coil after reducing *B*_*z*_ from approximately -40 to 0 *μ*T or reversing the magnetic field to +40 *μ*T to assess whether worms were responding primarily to gravity or instead to vertically skewed static magnetic fields (***figure Supplement 2***; see Methods). Under these conditions, we observed a strong upward migratory preference when the magnetic field was counteracted using a solenoid coil surrounding a Faraday cage (average vertical location = +2.56, n = 696 worms, 9 trials; p <0.0001, all comparisons). In an even more convincing experiment, we found that fully reversing the magnetic field from -40 *μ*T to +40 *μ*T also did not substantially alter the tendency of the animals to move upward (average vertical location = +2.17, n = 1,112 worms, 9 trials; p <0.0001, all comparisons, respectively). Thus, a static magnetic field cannot explain the tendency of the animals to migrate upward in the assay chambers.

Supplementing the solenoid coil assay, we also performed experiments in which the chambers were rotated to align them either parallel or perpendicular to the vector of Earth’s magnetic field as it entered the Faraday cage. (The direction and strengths of the geomagnetic field were determined using the Physics Toolbox Suite App for Android and iOS; see Methods). If worms were primarily responding to Earth’s magnetic field, we would expect to see strong upward movement in the parallel, but not the perpendicular assay, as the effect of the field would be negated in the latter condition. In both cases, however, we found a clear preference for upward migration (average vertical location = +3.94, n = 470 worms, 6 trials and +4.20, n = 400 worms, 5 trials, parallel and perpendicular assays, respectively; p <0.0001, all comparisons) (***figure Supplement 2***). Thus, the orientation of the assay either along or perpendicular to the field was irrelevant to the movement of the animals away from the Earth’s center.

Evidence for magnetotaxis in *C. elegans* was previously supported by the observation that AB1 wild isolate worms, which are native to Adelaide, Australia in the Southern Hemisphere, showed the opposite response to magnetic fields than N2, a strain isolated in Bristol, England ***Vidal-Gadea et al. (2015***) in the Northern Hemisphere. As the inclination of Earth’s magnetic field in Bristol is oriented approximately 180° relative to the field direction in Adelaide (66°and -66°, respectively; ***figure Supplement 4***), the investigators suggested that the opposing behavior of these strains could be explained by adaptation to their respective ambient electromagnetic fields. To further assess whether magnetosensation might guide vertical preference in our assay, we performed experiments comparing the movement of N2 and AB1 dauer larvae. We found that, similar to N2 dauers, AB1 dauers showed significant net upward migration compared with normally distributed data, albeit with a somewhat reduced bias (average vertical location = +2.44, n = 775 worms, 8 trials; p <0.0001, all comparisons; ***figure Supplement 4***). While this difference in the strength of vertical preference may suggest other interesting variations between N2 and AB1 worm behavior, it contradicts the conclusion that upwards migration in this assay is primarily driven by magnetotaxis, as has been described previously.

Each of these findings complement results from other labs, which have not found behavioral responses to magnetic field direction or intensity ***Landler et al. (2018a***); ***Njus et al. (2015***), except in instances in which worms have ingested paramagnetic particles ***Gourgou et al. (2023***). Moreover, in examining the data from one of these previously published experiments that tested vertical migration behavior within a solenoid coil, we noted an unreported trend in vertical migration behavior that is suggestive of gravitaxis behavior (***figure Supplement 5***, data replotted in new format and reprinted with permission). While these data were collected over a shorter distance under various conditions and analyzed by the index method (counting the number of worms migrating up vs down), comparison of the mean GI to a hypothetical mean of 0 revealed a statistical vertical preference in one instance when using a two-sided t-test (GI = 0.17+/- 0.08, SEM; p <0.05); however, it should be noted that this comparison was not found to be significant using the Wilcoxon signed-rank test, as was reported by the authors of that study. Together, these data collected across different laboratory settings point to a behavioral response that is consistent with gravitactic but not magnetotactic migration preference.

One concern that may be raised when assaying worms in a solenoid coil or along a large vertical distance is that their behavior may be influenced by thermal gradients along the vertical axis. Indeed, when measuring the temperature at the top and bottom of our Faraday cage, we found a consistent difference of ∼1°C as a result of rising warm air in the laboratory. This difference corresponds to a gradient of <0.02°C/cm, which is substantially below the reported 0.1°C/cm threshold temperature gradient to which worms are known to be sensitive ***Ramot et al. (2008***). However, as the reported threshold is limited by the accuracy of temperature measurements, worms may be capable of detecting even shallower gradients. To clarify whether temperature preference might be responsible for the vertical migration, we reversed the temperature differential in our Faraday cage, such that the bottom of the cage was 1°C warmer than the top by providing a heat source at the bottom (see Methods). Under this condition, we found that the pronounced upward migration persisted (average vertical location = +1.95, n = 445 worms, 5 trials; p <0.0001, all comparisons), although the average vertical movement was significantly lower than in the unaltered condition (p <0.0001; ***figure Supplement 6***).

Our findings indicate that the net vertical migration is not dramatically influenced by either magneto- or thermo-sensation. The pronounced upward migration of the animals was consistently maintained when the animals were shielded from light and alternating EMF, exposed to non-alternating magnetic fields in either of opposing directions, and subjected to weak temperature gradients in either orientation along the axis of the assay system. As there are no other obvious constant environmental influences that the animal is likely to experience that are oriented along the vertical axis besides the direction of Earth’s gravitational force under these various conditions, our findings provide strong support for the notion that it is the vector of the gravitational field that is responsible for the behavior, which we will henceforth refer to as negative gravitaxis ***Figure 2C–D***.

### Elimination of the neural-expressed MEC-7/12 microtubule subunits and MEC-5 collagen, but not other components involved in gentle touch sensation, abolishes negative gravitaxis

Gravitational force is exceedingly weak compared with other mechanical stimuli, and particularly so for organisms with low mass. For a dauer worm of roughly 150 ng in mass, based on size and reported estimates of overall density ***Kim et al. (2022***); ***Reina et al. (2013***), the Earth’s gravity would generate a maximum force of approximately 1.5 nN (see Methods for calculations). By contrast, worms are sensitive to a gentle touch input of 1-10 *μ*N ***Nekimken et al. (2020***); ***Petzold et al. (2013***). Diminutive organisms must therefore evolve extremely sensitive mechanisms for perceiving this minute force. In some small invertebrates, stretch receptors in the cuticle generate an ion influx upon deformation in a manner analogous to stretch and pressure sensors in mammals ***Bender and Frye (2009***). The comprehensive set of molecular components that mediate gentle touch sensation in *C. elegans* have been identified through extensive genetic analysis ***Gu et al. (1996***). Because the gentle touch mechanosensory channel proteins MEC-4 and MEC-10 are also essential for transducing hypergravitational force in worms ***Kim et al. (2007***), we sought to determine whether these known mechanosensory components function in sensing Earth’s gravity.

We found that mutations that eliminate most components involved in the gentle touch response ***Figure 3*** did not prevent gravitaxis in dauer larvae, demonstrating that gravity sensing is separable from touch reception. Gentle touch in *C. elegans* is perceived by the mechanosensory DEG/ENaC channels MEC-4 and MEC-10 ***Chalfie and Sulston (1981***), which form a pressure-sensitive channel in the set of Touch Receptor Neurons (TRNs). We found that neither *mec-4(-)* nor *mec-10(-)* mutants were significantly defective for gravitaxis compared with wildtype controls ***Figure 4***: *mec-4(e1339)* and *mec-10(tm1552)* mutant strains both exhibited a strong upwards directional bias in vertically oriented chambers compared to the horizontal controls and simulated normal distributions (average vertical location = +3.05, n = 1,340 worms, 7 trials and +2.94, n = 2,119 worms, 14 trials, respectively. p <0.0001, all comparisons) ***Figure 4***. An additional mutant strain of *mec-10* gave similar results ***figure Supplement 1***. Analysis of *mec-4(-)* mutants is complicated by the observation that *mec-4(tu253)* null mutant animals are sluggish ***Maguire et al. (2011***); this impaired movement could confound conclusions about the requirement for MEC-4 in gravitaxis. Indeed, when tested in our assay, *mec-4(tu253)* worms showed severely defective movement and did not show clear gravitaxis ***figure Supplement 1***. To rule out possible redundancy from either MEC-4 or MEC-10, we created and assayed a *mec-4(e1339); mec-10(tm1552)* double mutant strain, which similarly demonstrated a strong upward bias in our assay (average vertical location = +2.95, n = 481 worms, 6 trials; p <0.0001, all comparisons). Together, our findings with the *mec-4(e1339)* missense mutant, which is touch-defective, and the *mec-10(tm1552)* null mutant suggest that the MEC-4/10 channel are not required for negative gravitaxis.

**Figure 3.**
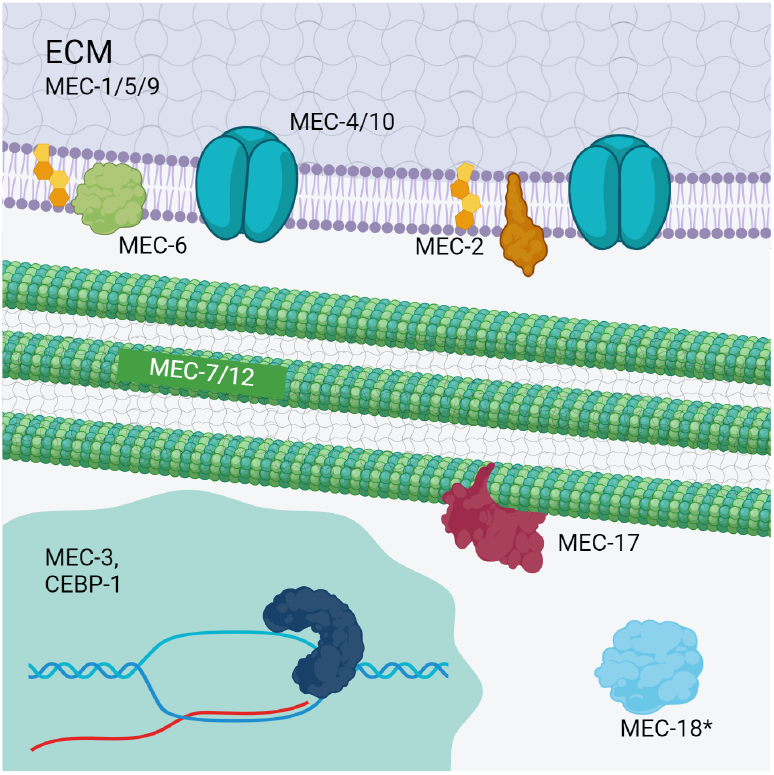
MEC-7/12 microtubule subunits, but not several other components in the gentle touch mechanosensory system, are necessary for gravitaxis. MEC-4 and MEC-10 subunits form a sodium channel that is required for gentle touch sensation in TRNs. MEC-1, -5, and -9 components of the extracellular matrix (ECM) form a structure between TRNs and the cuticle ***Cueva et al. (2007***); ***Emtage et al. (2004***). *—* MEC-7/12 microtubules form 15 protofilament structures in the TRNs only; however, these *β* and *α* subunits (respectively) are found in other neuronal 11 protofilament structures ***Bounoutas et al. (2009***). MEC-2 proteins associate with both the Deg/EnaC channels and TRN microtubules and enhance channel activity, likely through interactions with cholesterol in the plasma membrane ***Brown et al. (2008***); ***Chen et al. (2015***); ***Goodman and Schwarz (2003***). MEC-17 is a transacetylase that stabilizes MEC-7/12 microtubules ***Neumann and Hilliard (2014***); ***Shida et al. (2010***); ***Topalidou et al. (2012***). MEC-18 (starred) performs a poorly understood role the transduction of gentle touch. Based on its proposed function as an acetyl CoA ligase, we propose it may act in conjunction with MEC-17 to acetylate tubulin proteins. (Adapted from images by ***Tavernarakis and and Driscoll (1997***) and others). Image was created with BioRender.com.

**Figure 4.**
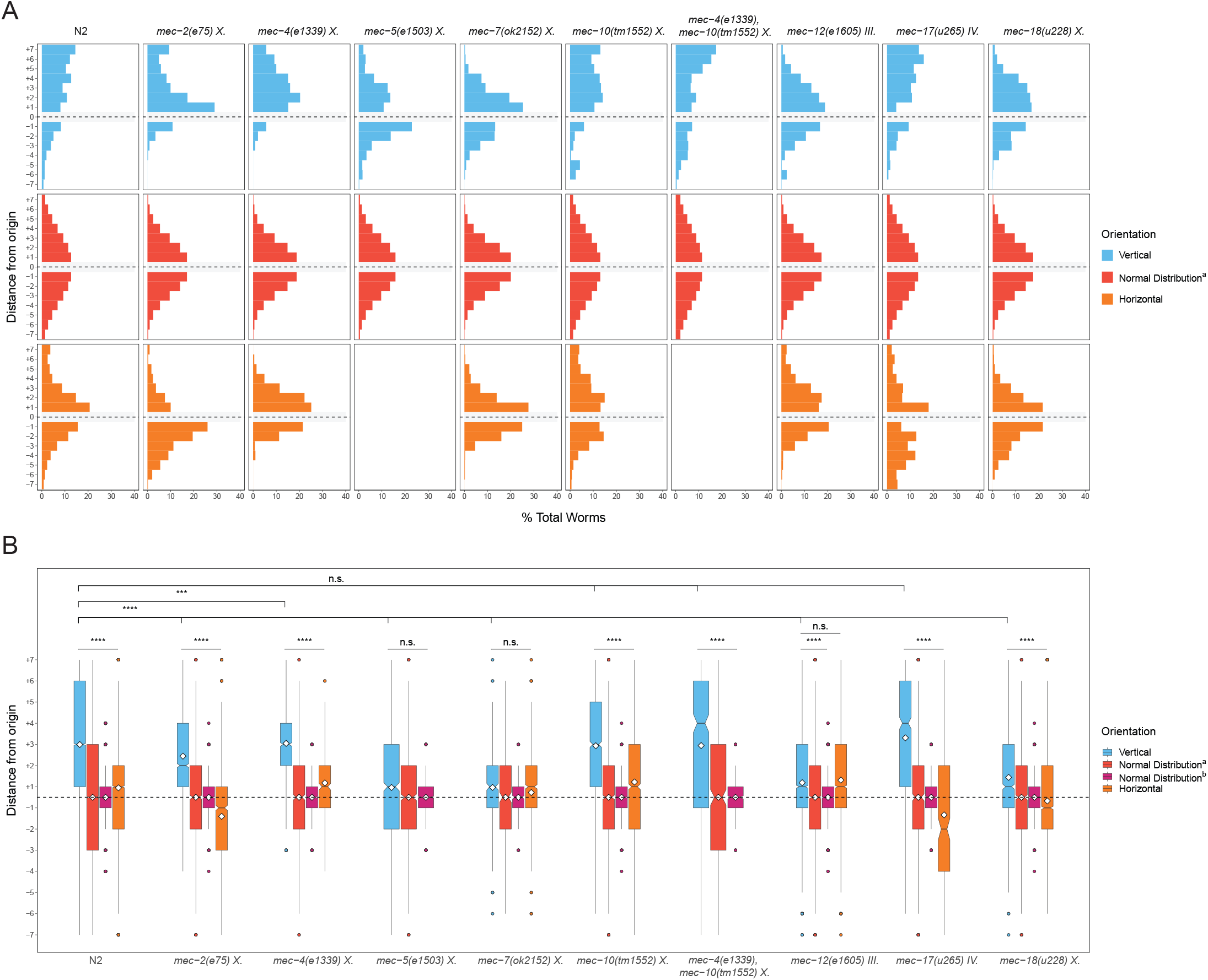
*C. elegans* negatively gravitax under different environmental conditions. **(A–B)** Gravitaxis assays of N2 dauers and dauers carrying mutations in genes involved in mechanosensory transduction. **A** Histograms depicting cumulative distribution of *C. elegans* worms over multiple trials in vertical (blue) and horizontal (orange) assays as well as in comparison with a simulated normal distribution (Normal Distributionv), which is created using a mean of 0 and the same standard deviation as the combined vertical assay distributions. **B** Boxplots summarizing data shown in A. * p <0.05, ** p <0.01, *** p <0.001, **** p <0.0001; n.s. is not significant using Kruskal-Wallis followed by Dunn’s test with Bonferroni correction. Notches on boxplots represent 95% confidence intervals; mean values are indicated with a diamond. Normal Distribution^*a*^ is described above; Normal Distribution^*b*^ was created using a mean of 0 and the same sample size as the combined vertical assays. Dotted horizontal line indicates the origin of the assay (location “0”), which was not scored. **Figure 4—figure supplement 1**. Gravitaxis behavior in additional strains **Figure 4—figure supplement 2**. Worm distribution variation between trials

We found that several other genes in the canonical gentle touch sensing system are similarly largely dispensable for gravitaxis. *mec-2* encodes a stomatin-like protein required for MEC-4/MEC-10 channel activity ***Brown et al. (2008***). MEC-2 binds cholesterol ***Huber et al. (2006***) and likely facilitates gentle touch transduction by altering the composition of the plasma membrane surrounding MEC-4/MEC-10 ion channels ***Brown et al. (2008***). We found that *mec-2(e75)* dauers were not significantly impaired in their ability to gravitax when compared with the horizontal controls and normal distributions (average vertical location = +2.45, n = 2,621 worms, 10 trials; p <0.0001, all comparisons; ***Figure 4***). MEC-18 is required for gentle touch sensation, although its exact function has not been well characterized ***Gu et al. (1996***); ***Zhang and Chalfie (2002***). *mec-18(u228)* mutants showed a strong gravitactic preference compared to controls (+1.44, n = 1,645 worms, 4 trials; p <0.0001, all comparisons). While both mutants showed a significantly reduced vertical preference compared to N2 vertical distributions (p <0.0001 in both cases), these experiments point to at most a minor role for these components in gravitaxis. In contrast, disruption to MEC-5, a collagen protein associated with the gentle touch transduction pathway, resulted in a notable decrease in gravitactic behavior (average vertical location = +0.96, n = 1,051 worms, 6 trials; p >0.05, all comparisons; ***Figure 4***). Though research on MEC-5 in mechanosensation has largely focused on TRN/gentle touch transduction, MEC-5 is also expressed in ciliated head neurons ***Hammarlund et al. (2018***); ***Taylor et al. (2020***) and is essential for male mating behaviors through CEM neurons ***Vore et al. (2018***), suggesting a potential role for ciliated neurons in gravity transduction. We also found that a missense mutation in *mec-6* disrupted gravitactic behavior (average vertical location = +0.26, n = 195 worms, 6 trials; p >>0.05, all comparisons). However, as with *mec-4* null worms, these animals are known to be sluggish, consistent with their limited migration and narrow distribution around the origin of our assay ***figure Supplement 1***.

The specialized MEC-12 alpha-tubulin and MEC-7 beta-tubulin proteins form unique 15-protofilament microtubules that are found only in the six TRNs and that are essential for mechanosensation. TRN microtubule bundles are thicker than the typical 11 protofilament microtubules found in most cells, including in other neurons ***Chalfie and Thomson (1982***). These 15 protofilament structures may provide intracellular resistance required for mechanotransduction; however, their exact function in this process is unknown ***Bounoutas et al. (2009***); ***Krieg et al. (2015***). In contrast to our findings with other gentle touch sensation components, we found that removal of either MEC-7 or MEC-12 abolished negative gravitaxis, resulting in a random distribution in the chambers similar to that seen in the horizontal controls, although *mec-12* mutant strains differed significantly from simulated normal distributions (average vertical location = +0.97, n = 407, 4 trials; p >>0.05, all comparisons for *mec-7(ok2152)* and average vertical location = +1.19, n = 5,099, 12 trials; p >>0.05, horizontal comparison and p <0.0001 comparing against normal distributions for *mec-12(e1605)*; ***Figure 4***). A strain containing a missense mutation in *mec-7* similarly demonstrated little to no gravitactic preference ***figure Supplement 1***.

MEC-7 and MEC-12 are required for normal axonal outgrowth ***Bounoutas et al. (2009***); ***Chalfie and Thomson (1982***) and it was therefore conceivable that the *mec-7(-)* and *mec-12(-)* mutations might block gravitaxis primarily by altering neuronal development or structure. We were able to separate the role of MEC-7/12 in neuronal structure from its other functions by taking advantage of the *mec-12(e1605)* allele, an H192Y missense mutation that eliminates gentle touch sensation without detectably altering TRN development or structure ***Chen et al. (2014***); ***Lockhead et al. (2016***). We found that gravitaxis is greatly diminished in *mec-12(e1605)* mutants Fig3B,C, suggesting that MEC-12 requirement in gravitaxis is separable from its role in neuronal morphology.

MEC-12 is the only *C. elegans* alpha-tubulin subunit known to be acetylated at K40 ***Akella et al. (2010***); ***Lockhead et al. (2016***). Mutations that modify K40 or that eliminate the MEC-17 transacetylase required for microtubule acetylation also prevent mechanosensation ***Neumann and Hilliard (2014***); ***Shida et al. (2010***); ***Topalidou et al. (2012***). We found that *mec-17(u265)* mutants, which lack functional MEC-17, undergo normal gravitaxis behavior (average vertical location = +3.31, n = 856 worms, 5 trials p <0.0001, all comparisons; ***Figure 4***) that did not significantly differ (p >>0.05) from that of N2 animals, indicating that this modification, which is essential for stabilizing the MEC-7/12 microtubules in TRNs, is not required for sensation or response to gravity.

Taken together, these results support the notion that MEC-7/12 microtubules are essential for gravity perception in a role that is distinct from their structural roles or action in conferring gentle touch sensitivity.

### The TRPA-1 channel is essential for gravity sensation

Our finding that MEC-4 and MEC-10 are not required for gravitaxis suggests that other types of mechanosensory channels might instead participate in gravity perception. Prime candidates for such channels are members of the superfamily of TRP (transient receptor potential) proteins, which are implicated in many sensory modalities, including sensitivity to touch, hot and cold temperatures, noxious chemicals, and light ***Chatzigeorgiou et al. (2010***); ***Venkatachalam et al. (2014***). Orthologs of these channels have been found across metazoan phylogeny, including in all triploblast and diploblast animals, sponges, and even unicellular choanoflagellates, which are believed to be the closest surviving relatives of all metazoans ***Carr et al. (2008***). Mouse TRPA-1 is expressed in the vestibular system, the primary organ where gravity sensation occurs in mammals ***Kamakura et al. (2013***). In *C. elegans, trpa-1* is expressed in the proprioceptive/nociceptive PVDL/R neuron pair, among others. Although TRPA-1 is known to play a role in noxious cold sensation in these neurons, *Drosophila* homologs of TRPA channels, *pain* and *pyx*, are required for gravity sensation ***Sun et al. (2009***). Additionally, *trpa-1* knockout mice show impaired mechanosensation and perception of noxious cold ***Kindt et al. (2007***). *C. elegans* TRPA-1 is likely to perform mechanosensory functions in addition to its known role in cold sensation ***Kindt et al. (2007***).

We found that removal of TRPA-1 in the *trpa-1(ok999)* null mutant abolishes gravitaxis in dauer larvae (average vertical location = +1.36, n = 1,392, 10 trials; p >>0.05, all comparisons; ***Figure 5A–B***). As *trpa-1(ok999)* mutant adult worms exhibit several movement defects, including decreased forward locomotion and several variations in the sinusoidal movement typical of wildtype worms ***Yemini et al. (2013***), it was conceivable that elimination of net upward biased movement of the animals might reflect diminished locomotory capacity rather than defects in gravity sensing per se. However, we found that *trpa-1(ok999)* dauer larvae distributed broadly across the chambers comparable to that observed with N2 animals in both the horizontal and vertical conditions, demonstrating that these mutants are capable of traveling long distances regardless of orientation. These findings suggest that TRPA-1 channels are essential for, and may mediate, *C. elegans* gravitaxis.

**Figure 5.**
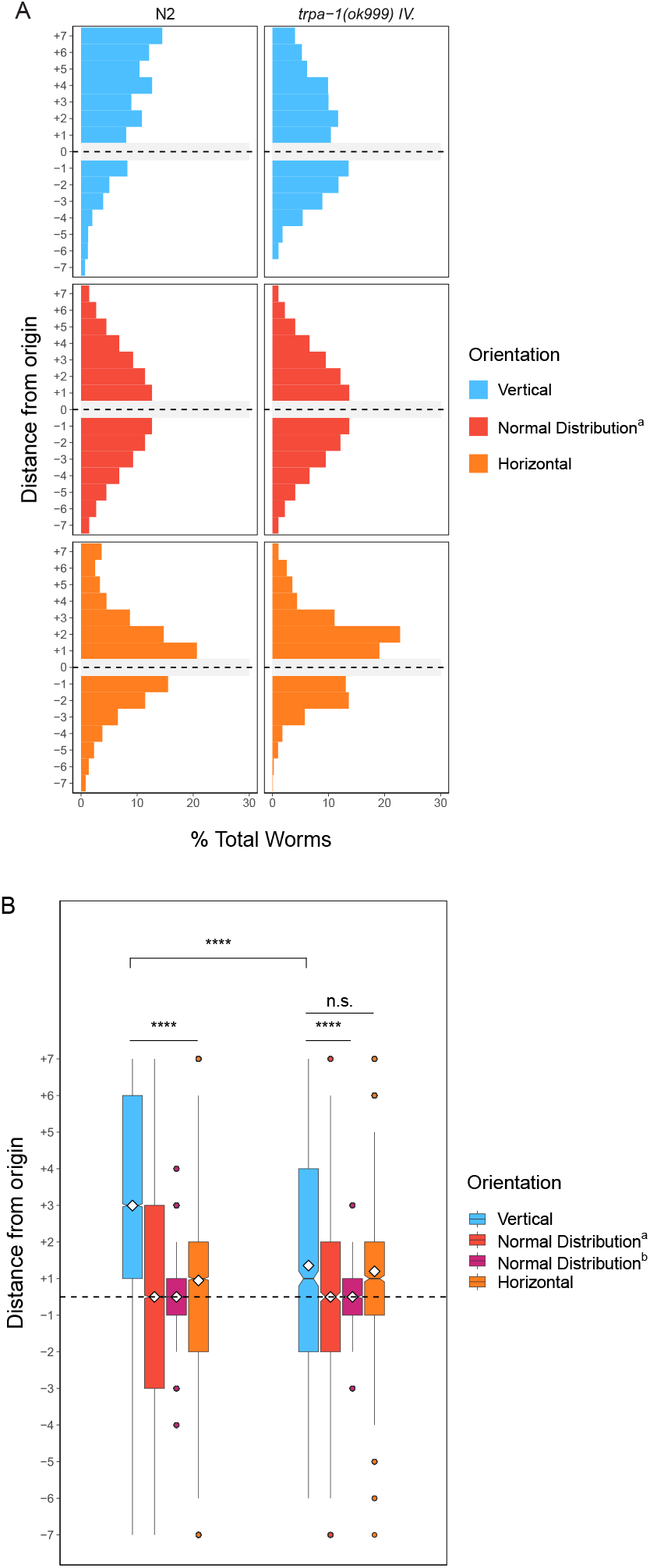
TRPA-1 channels are required for gravitaxis. **(A–B)** *trpa-1* mutants in vertical (blue) and horizontal (assays) compared with N2 controls, as well as simulated normal distributions (described in previous figure legends and Methods). ∗ p <0.05, ∗∗ p <0.01, ∗∗∗ p <0.001, ∗∗∗∗ p <0.0001; n.s. is not significant using Kruskal-Wallis followed by Dunn’s test with Bonferroni correction. Notches on boxplots represent 95% confidence intervals; mean values are indicated with a diamond.

### PVD neurons, but not TRNs, are required for gravitaxis

The requirement in gravitaxis for *trpa-1, mec-5, mec-7*, and *mec-12*, but not *mec-4/10* or other genes required for gentle touch response suggested that neurons other than the TRNs are required for this behavior. Although MEC-7 and MEC-12 are present in many neurons, their function beyond the TRNs is poorly understood. Further, there is no reported connection between MEC-7/12 microtubules and TRPA-1 function. Of the six TRNs, only one expresses *trpa-1* ***Taylor et al. (2020***); however, available expression data ***Chatzigeorgiou et al. (2010***); ***Taylor et al. (2020***) indicate that PVDs, which detect muscle movement in addition to a host of other stimuli, express all three genes. While OLL and AFD neurons also co-express MEC-7/12 and TRPA-1 ***Hammarlund et al. (2018***); ***Taylor et al. (2020***), the proprioceptive role of PVD neurons led us to hypothesize that PVD neurons might be integration centers for two proprioceptive inputs: muscle movement and position relative to the vector of Earth’s gravitational field.

To test whether PVD neurons are important for gravitaxis, we first investigated the requirement for TRNs in gravity sensing by genetically ablating them. The *mec-4(e1611d)* gain-of-function mutation results in constitutive MEC-4/10 activation, which triggers degeneration and necrotic death of the TRNs specifically. Ablation of the TRNs did not alter the ability of the animals to perceive and respond to gravity: the gravitactic behavior of worms lacking TRNs was not significantly different from that of N2 animals in our gravitaxis assay (p >>0.05) (average vertical location = +2.92, n = 1,955, 12 trials; p <0.0001, all comparisons; ***Figure 6A–B***); hence, efficient gravitaxis of dauer larvae does not require the TRNs.

**Figure 6.**
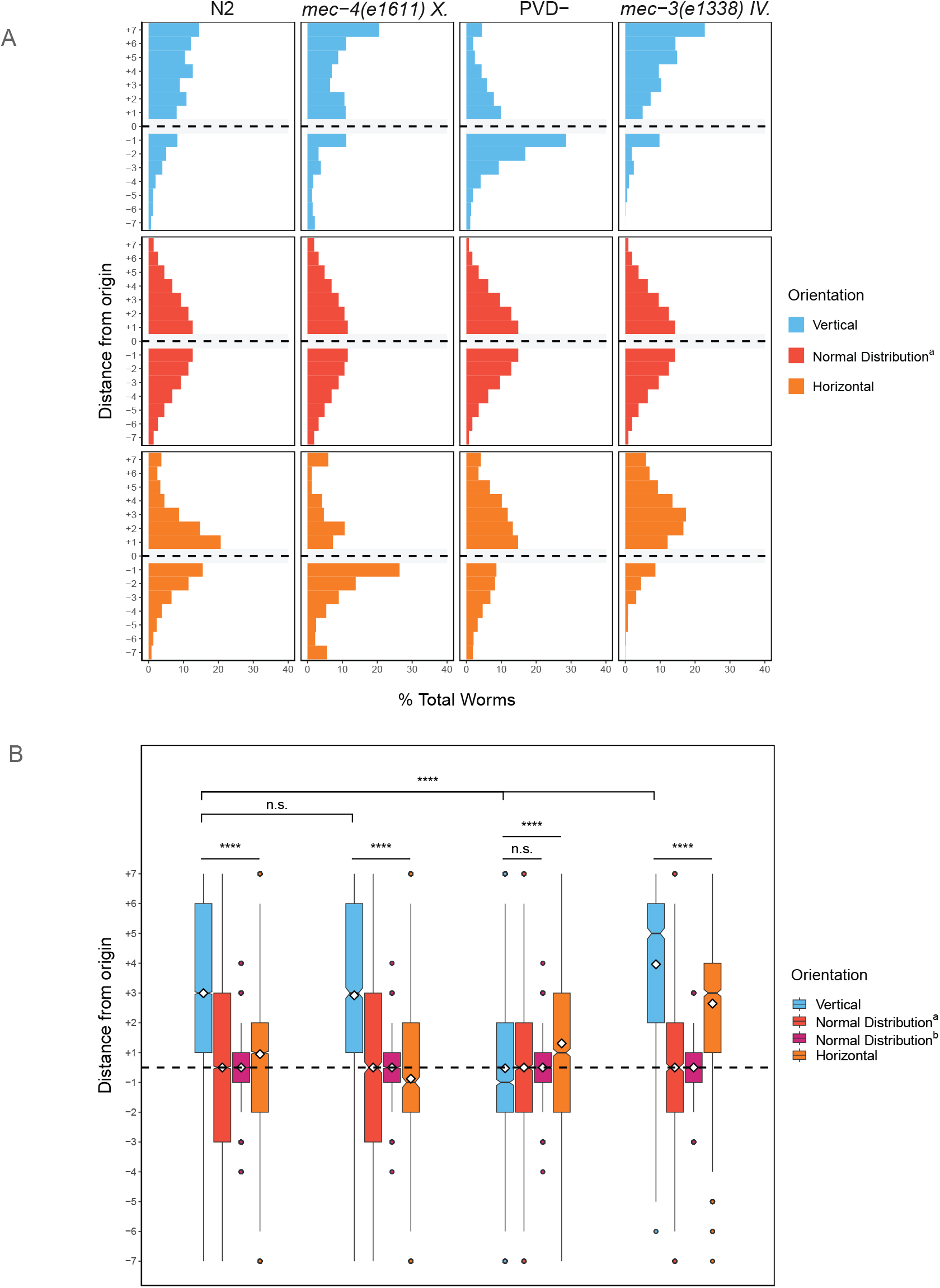
Genetic ablation of PVDs but not TRNs affects gravitaxis. **(A–B)** Gravitaxis behavior of *mec-3(e1338)* dauers, dauers lacking TRNs only (*mec-4* gof), dauers lacking PVD neurons only (*ser2prom-3::deg-3*), and N2 dauer controls. * p <0.05, ** p <0.01, *** p <0.001, **** p <0.0001; n.s. is not significant using Kruskal-Wallis followed by Dunn’s test with Bonferroni correction. Notches on boxplots represent 95% confidence intervals; mean values are indicated with a diamond.

We then ablated the PVD neurons by expressing a constitutively active version of the nicotinic acetylcholine receptor (nAChR) channel subunit, *deg-3(u662)*/DEG-3, under the *ser2prom3* promoter ***Albeg et al. (2011***). Elimination of PVD neurons using this construct has been confirmed with a PVD-specific fluorescent reporter ***Albeg et al. (2011***). (Importantly, this construct was not found to ablate OLL neurons, which also show expression from the *ser2prom3* promoter.) We found that animals lacking PVD neurons completely failed to undergo negative gravitaxis, showing a relatively even distribution between upward and downward movement (average vertical location = +0.47, n = 2,348 worms, 10 trials; p <0.0001, all comparisons; ***Figure 6A–B***). This behavior is likely attributable to *bona fide* inability of the PVD-ablated animals to undergo negative gravitaxis, rather than defects in motility, as they are capable of effectively reaching both ends of the chamber in either the horizontal or vertical orientations. These findings strongly support an essential requirement for PVD neurons in sensing and responding to gravitational fields.

Our understanding of the role of PVD, TRN, and other sensory neuron involvement in gravitational response is complicated by the fact that dauers, which undergo arrested development, have distinctive neural architecture compared with adults. For example, PVD neurons in dauers lack 4° processes that are involved in communicating proprioceptive feedback to muscle cells ***Tao et al. (2019***) ***Figure 7***. Given that PVD neurons in particular are highly compartmentalized such that harsh touch and proprioceptive signaling occur within discrete parts of the cell (i.e., 1° axons and 2°-4° dendritic processes)l ***Tao et al. (2019***), we hypothesized that some of these processes may be dispensable in regulating gravitactic behavior. To examine this question, we assayed the gravitactic preference of *mec-3(e1338)* mutant worms, in which a frameshift mutation alters the function of the LIM homeodomain transcription factor MEC-3. MEC-3 mediates differentiation of several sensory neurons including PVDs and TRNs by activating a battery of genes required for touch sensation. TRNs in mec-3(-) mutants exhibit smaller-than-normal cell bodies and shorter processes, leading to touch insensitivity ***Chalfie and Sulston (1981***), while PVD neurons lack 2°-4° processes in adults ***Smith et al. (2010***); ***Tsalik et al. (2003***). We found that elimination of MEC-3 function in a *mec-3(e1338)* mutant did not significantly alter the ability of the animals to undergo gravitaxis (average vertical location = +3.96, n = 1,225 worms, 5 trials; p <0.0001; ***Figure 6***). Thus, normal TRN and PVD differentiation is apparently not essential for negative gravitaxis. The requirement for PVD neurons in this response may be limited to the development of their cell bodies and 1° axonal processes.

**Figure 7.**
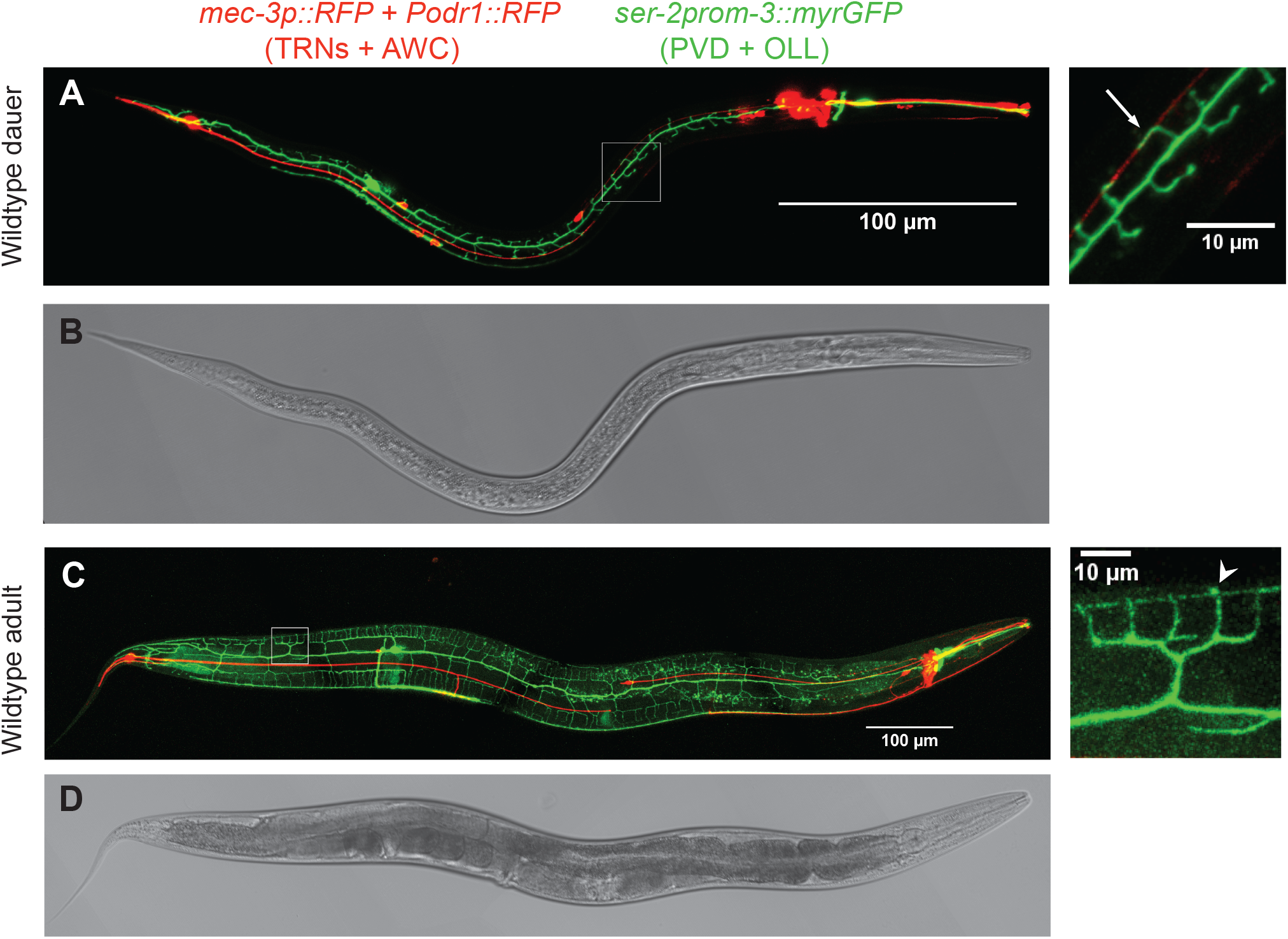
PVDs and TRNs in dauer and adult worms. **(A and C)** Confocal fluorescent micrographs of *mec-3p::RFP* and *ser-2prom-3::GFP* expression labelling the TRN and AWC neurons and PVD and OLL neurons, respectively. Brightfield images shown in **B** and **D. (A-B)** Wildtype dauer worms; developing 3° processes of PVD neurons are indicated with a white arrow in inset. **(C-D)** Wildtype adult hermaphrodites; 4° process are indicated with white arrowhead in inset.

The difference in PVD architecture between dauers and adults led us to ask whether adults behave differently in response to gravity. Changes in gravitational response – or even in the ability to sense gravity – throughout development could also be important for the worms’ ecology, particularly given the importance of the dauer stage, but not adults, in dispersal of the animal. We found that, like dauer larvae, N2 adults also undergo significant negative gravitaxis in the absence of light and EMF (average vertical location = +2.36, n = 548, 8 trials; p <0.0001; ***Figure 8A–B***). While overall gravitactic preference was reduced somewhat in adults compared with dauers in our vertical assay (p <0.01), we note that the distances travelled by adults and dauers may not be directly comparable as adults are significantly larger, and therefore travel shorter distances relative to their body length in this assay. Evidence of gravitactic behavior by adults suggests that gravity sensation may have a strong influence on behavior not only in the dispersal stage, but throughout *C. elegans* development.

**Figure 8.**
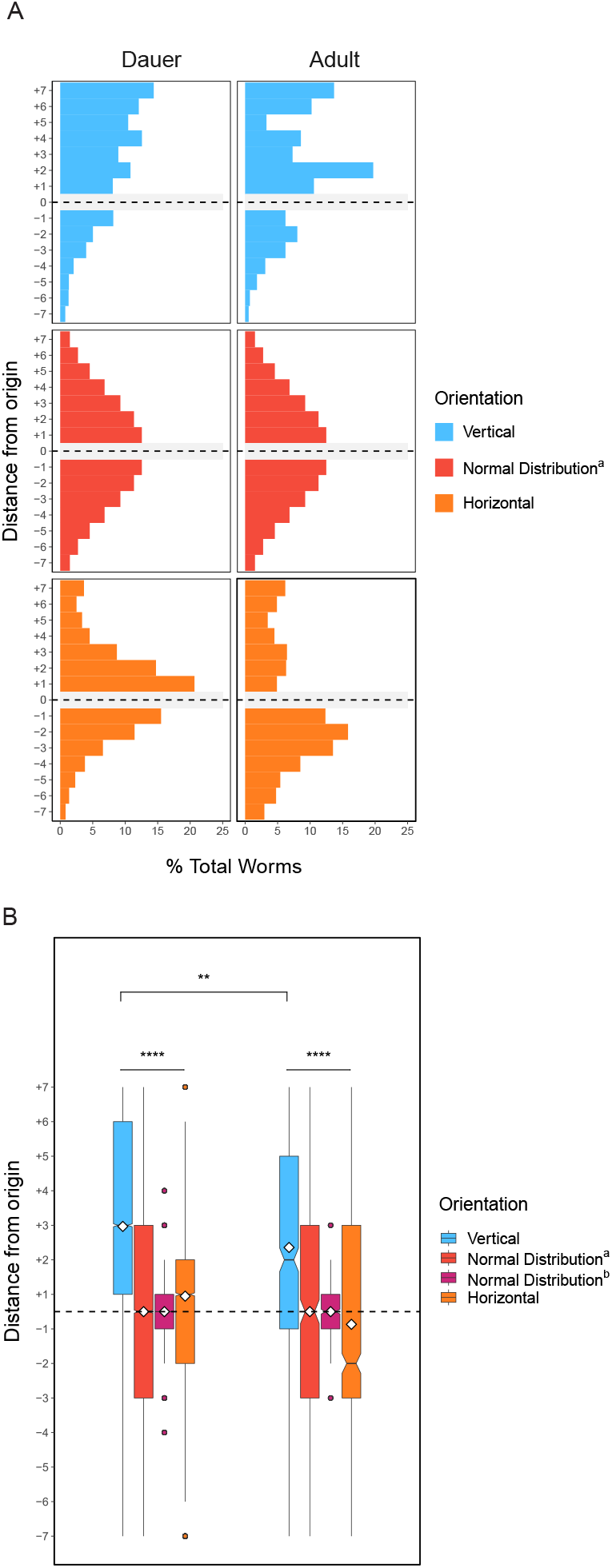
Adult worms negatively gravitax. **(A–B)** Adult worms demonstrate a strong negative gravitaxis preference similar to dauers. ∗ p <0.05, ∗∗ p <0.01, ∗∗∗ p <0.001, ∗∗∗∗ p <0.0001; n.s. is not significant using Kruskal-Wallis followed by Dunn’s test with Bonferroni correction. Notches on boxplots represent 95% confidence intervals; mean values are indicated with a diamond.

## Discussion

In this study, we report four major advances. 1) We discovered that *C. elegans* dauer larvae and adults show pronounced negative gravitaxis. This drive toward movement away from the center of the earth may direct the animals toward food sources, typically decomposing vegetative matter at the surface ***Frézal and Félix (2015***). 2) We found that the ability of the animals to sense and respond to the force of gravity is attenuated by light and electromagnetic fields, suggesting that they integrate sensation of these other environmental influences to make decisions about whether to undergo negative gravitaxis. 3) The ability of the animals to sense and/or respond to gravity does not require DEG/ENaC channels or other factors involved in touch sensation but does require the MEC-7/12 microtubule components, MEC-5 collagen, and TRPA-1 channels. Thus, gravity sensing involves a previously unknown system involving specialized microtubules, which also participate in touch sensation and may function with TRP sensory channels. 4) Both immature PVD neurons in dauer larvae, and those in adults, but not TRNs, are essential for response to gravity. Moreover, adult PVDs lacking 2°-4° processes (“menorahs”) are fully functional in the gravitaxis assay, demonstrating that these elaborations are not required for response to gravity.

Our findings that *C. elegans* dauer larvae exhibit pronounced negative gravitaxis contrasts with prior studies which reported no gravitactic behavior in dauers, or that in which positive reorientation toward the gravitational vector in adults in a liquid suspension was observed ***Chen et al. (2021***); ***Okumura et al. (2013***). These differences are likely attributable to differences in several parameters of assay design. Previous experiments have been performed on Petri dishes, which limit the distance animals can travel. Owing to the long distances traversed in our chambers, we were able to find robust and highly significant differences between vertically oriented chambers and normal distributions or paired horizontal controls. In addition, the environment of a worm crawling in the thin space between the agar substrate and the wall of the chamber may be more similar to the act of crawling through a column of soil than the experience of moving along the agar surface at the interface with the atmosphere on a plate. Finally, earlier experiments on gravitaxis in *C. elegans* made no mention of the lighting conditions used or shielding against EMF, leading us to propose that signal integration with gravity and these other sensory inputs may have attenuated the behavior in those studies, as we have demonstrated here.

Underground habitats are generally ill-suited for *C. elegans*. Such environments may cause dauer larvae to find their way to the surface where decaying vegetation might be present. However, the animals also avoid light ***Ward et al. (2008***), which is associated with damaging ultraviolet radiation. Some studies suggest that the animals may navigate using the Earth’s magnetic field ***Landler et al. (2018a***); ***Vidal-Gadea et al. (2015***, 2018), although this finding has been greatly debated ***Landler et al. (2018a***,b); ***Malkemper et al. (2023***); ***Njus et al. (2015***); ***Vidal-Gadea et al. (2018***). Our finding that ambient light and EMF in the laboratory setting strongly attenuate gravitactic behavior – likely explaining why negative gravitaxis had not been reported previously – suggests that the animals integrate sensory inputs from a variety of environmental signals, including the force of gravity, to optimize behavioral decisions ***Metaxakis et al. (2018***) in complex environments. Additionally, by observing gravitaxis behavior under conditions in which the weak effects of Earth’s magnetic field and shallow thermal gradients are further minimized, we found that gravity, and not magnetic or temperature stimuli, is the only obvious influence that leads to upward migration. The extent to which light, EM fields, and potentially other inputs override the response to gravity, as well as other sensory neurons and the circuits required to compute their respective influence on behavior, are important outstanding questions.

Although it was not detected in earlier studies, our altered assay system, which effectively shields the animals from other confounding environmental stimuli, revealed that both dauers and adults of *C. elegans* exhibit pronounced negative gravitaxis. The reduced ability of N2 dauers to undergo negative gravitaxis compared to that observed with *C. japonica* dauers might be an outcome of myriad adaptations to laboratory cultivation, as has been observed for several other traits, including social feeding, egg laying behavior, oxygen tolerance, and nictation ***de Bono and Bargmann (1998***); ***Félix and Braendle (2010***); ***Frézal and Félix (2015***); ***Gray et al. (2004***); ***Large et al. (2017***); ***Lee et al. (2012***). We found that negative gravitactic behavior, albeit weakened, is also seen with the LSJ1 sister strain ***figure Supplement 1***, which has been raised in liquid media over many decades, as well as in *C. briggsae* ***Ackley et al. (2022***). It remains an open question as to whether these strains show negative gravitaxis when shielded from light and EMF when swimming in liquid rather than crawling on solid media.

Our finding that TRPA-1, MEC-5 collagen, and MEC-7/12 microtubule functions are required for gravitaxis reveals an unexplored mechanism for transducing gravitational force. Unlike the DEG/ENaC channels, TRPA-1 receptors do not require internal or external machinery for their function ***Kindt et al. (2007***). Additionally, *trpa-1* is expressed in only one of six TRNs, which are the only neurons containing the 15 protofilament structures formed by MEC-7/MEC-12 tubulins. MEC-7/MEC-12 microtubules may perform more than a structural role in mechanosensation; whether this function is independent of their assembly into 15 protofilament structures has yet to be determined.

We obtained evidence pointing to a pair of neurons, PVDL/R, as neural components of the gravity-sensing system. Unlike animals lacking TRN function or cells, which showed normal behavior in our gravitaxis assay, those lacking PVD cell bodies and axons show no detectable response to gravity. These results, in conjunction with our *trpa-1* findings, suggest that TRPA-1 expression in PVD neurons may be important for this. However, these observations alone are insufficient to conclude that TRPA-1 is either the primary, or sole gravity receptor in worms. As with TRP channels generally, TRPA-1 is known to perform primary and secondary roles in a variety of sensory transduction pathways ***Montell (2003***). It is not clear from our studies, or other reports, exactly how MEC-7 and MEC-12 microtubule subunits or MEC-5 collagen contribute to mechanosensation. While tubulins may perform a vital function within PVD neurons that are required to transmit a signal, it is also possible that the structures play a role in other neurons involved in the response. It is noteworthy that ciliated neurons are frequently used in gravity sensation including in humans ***Bezares-Calderón et al. (2020***); ***Lacquaniti et al. (2014***). *C. elegans* contains 60 ciliated neurons ***Albeg et al. (2011***), 38 of which express both MEC-12 and TRPA-1 ***Hammarlund et al. (2018***); ***Taylor et al. (2020***) and some of which may require MEC-5 collagen in order to function properly ***Vore et al. (2018***). Of these 38 cells, 3 (OLQ, AQR, and PHA) also express an orthologue of the likely gravity-sensitive mechanoreceptor in mammals, TMC-1 ***Chatzigeorgiou et al. (2013***); ***Corey and Holt (2016***); ***Nist-Lund et al. (2019***). Thus, it is conceivable that PVDs may function as secondary neurons acting within a neural circuit to make decisions about movements in response to the direction of the gravitational field and other stimuli.

Understanding how *C. elegans* perceives gravity has implications for both invertebrate and mammalian biology. *C. elegans* has been used for decades as a model for understanding the effects of microgravity on animal physiology in space ***Gao et al. (2015***, 2017); ***Honda et al. (2012***); ***Qiao et al. (2013***); ***Selch et al. (2008***); ***Xu et al. (2014***); ***Zhao et al. (2017***), and yet scant information is available regarding their ability to sense gravity. Moreover, gravity perception is linked evolutionarily and developmentally with hearing ***Fritzsch et al. (2014***); ***Lipovsek and Wingate (2018***): the vestibular and auditory systems in mammals share similar sensory structures, transduction machinery, and gene expression ***Scheffer et al. (2015***). A role for TRPA1 has been identified in sensory transduction within inner ear hair cells in mice ***Corey et al. (2004***). The similarities between these systems make the study of gravity sensation important not only for understanding mammalian vestibular sensation, but also for auditory sensation. A recent study showed that *C. elegans* can sense sound through mechanical perturbations of the epidermis and that this sense involves the multimodal PVD neurons as well as other mechanosensory neurons ***Iliff et al. (2021***). Thus, uncovering the role of TRPA-1 as a mechanoreceptor raises the possibility that *C. elegans* could be used as a model for studying both vestibular sensation and audition. The warping of space-time caused by Earth’s mass is among the most consistent stimuli experienced by organisms across billions of years of evolution. Our results raise the exciting possibility that a mechanism for detecting this stimulus may have been preserved since at least the divergence of the nematode (*C. elegans*) and chordate (e.g., mammals) phyla over 500 million years ago.

## Methods and Materials

### Resource Availability

#### Lead Contact

Further information and requests for resources and reagents should be directed to and will be fulfilled by Joel Rothman.

#### Materials availability

Worm strains generated in this study will be made available upon request.

#### Data and code availability

All data reported in this paper will be shared by the lead contact upon request.

This paper does not report original code; scripts created for data analysis and visualization will be made available upon request.

Any additional information required to reanalyze the data reported in this paper is available from the lead contact upon request.

### Experimental Model and Subject Details

*C. elegans* worms were grown and maintained on OP50 *E*.*coli* at room temperature (23°C) using standard methods ***Stiernagle (2006***). Dauer formation was induced by overcrowding, which occurred 8-12 days after chunking onto seeded NGM plates. Adults used for gravitaxis assays and imaging were taken from synchronized populations 2 days after hatching. Crosses were performed as described previously ***Fay (2006***), using fluorescence microscopy, PCR, and/or sequencing to confirm the outcome of each cross.

All strains used in this study are cataloged in the Supplementary Material.

### Method Details

#### Dauer Isolation

Dauer larvae were isolated from 1-5 starved NGM plates using 1% SDS and a 30% sucrose gradient as described previously ***Karp (2018***); ***Ow and Hall (2015***). Briefly, worms were rinsed and collected in M9, then pelleted and resuspended in 7mL 1% SDS in a 15 mL falcon tube. Worms were left rotating for 30 minutes in SDS to kill all non-dauer life stages. After 3-5 rinses with M9, worms were resuspended in 10 mL 30% sucrose solution and centrifuged for 5 minutes. Surviving dauers were collected at the interface of the water-sucrose gradient with a large bore glass Pasteur pipette and rinsed 3-5X in M9. Dauers were used immediately or after rotating in solution overnight.

#### Synchronization

Mixed stage worms were rinsed from confluent plates with M9 and collected in 1.5 mL tubes. Worms were then rinsed and pelleted. A 1 mL bleach solution containing 150 *μ*L bleach and 50 *μ*L 5N KOH was added and worms were lysed in this solution for 3 minutes, or until most corpses were dissolved. The embryo pellet was then rinsed 3-5X in M9 and plated onto OP50 NGM plates until adulthood (∼2 days at 23°C).

#### Assay Preparation

Gravitaxis assay chambers were created using 5 mL serological pipettes and 4% NGM agar as described previously ***Ackley et al. (2022***). To assemble each chamber, a heated blade was used to cut the tapered end of one pipette and the cotton plug was removed from the end of a second pipette. The two pipettes were then joined end-to-end by melting the two ends slightly over a Bunsen burner and fusing them with moderate pressure, ensuring that no gaps remained. The NGM media was then autoclaved and then taken up by each double pipette using a standard serological pipettor while the agar was still molten. Best results were achieved by orienting each pipette parallel with the benchtop as much as possible during the procedure and as the agar solidified. Pipettes were used within 24 hours of construction or stored at 4°C to be used within 48 hrs.

Once solidified, a heated 3mm Allen key was used to punch a hole through one wall of the pipette about 5mm below the fusion point (for reference, the cotton-plug end is the “top”). Each end of the pipette was removed using a heated blade and sealed with parafilm.

#### Gravitaxis Assay

Worms were pelleted in M9 and were either extracted directly from the pellet or pipetted onto a square of parafilm. After cutting the tip to increase the bore, a pipette was used to aspirate 0.5-2 *μ*L of the concentrated worm pellet, letting in a small volume of air at the end to facilitate injection. Worms were injected into the agar at the opening created by the Allen key. This opening was then sealed with parafilm, and the chamber was immediately oriented vertically or horizontally. Each chamber represents an experimental trial in which worms were later scored individually. Separate conditions (such as different strains or environmental variables) were always run in parallel with at least one (but often multiple) N2/vertical/Faraday cage control.

Unless indicated otherwise, all experiments were performed in a custom-manufactured Faraday cage. The effectiveness of the cage in blocking EMF was tested by placing a cell phone within the cage and calling it to determine if it received a signal. Worms were allowed to gravitax overnight and were scored the following day (typically 12-24 hrs after injection).

Pipettes were examined under a dissecting microscope and a marker was used to mark the location of each worm. Pipettes were scored immediately after removal from the assay location. Worms were only scored if they appeared alive, healthy, and were not swimming (chambers were only scored if a majority of worms were alive and crawling). Worms were not scored within 2.5 cm to either side of the injection point.

To tally the location of each worm, pipettes were divided into 7 sections on each side of the 5 cm injection site. Each section was 3.5 cm long (the same length as a 1 mL volume indicated on the pipettes). The number of marker points in each section was then recorded. Data collection was validated with blinded replicates.

#### Solenoid coil, parallel vs. perpendicular, and temperature gradient experiments

A custom solenoid coil was constructed using a cardboard poster tube ∼60 cm in length and 10 cm diameter wrapped with 30 g wire spaced roughly 1 mm apart. For each assay, a Faraday cage consisting of aluminum foil and wire mesh was used inside the solenoid coil and a similarly constructed Faraday cage was used outside the coil in parallel. The Physics Toolbox Suite app for Android and iOS was used to measure the magnitude and direction of Earth’s magnetic field inside and outside the coil. The voltage from a Wanptek DC power supply unit (model number DPS305U) was then adjusted to reach an internal magnitude of 0 *μ*T along the z-axis inside of the solenoid coil. Experiments in the coil were run overnight as in other assays.

For parallel vs. perpendicular assays, chambers within our original Faraday cage were oriented with (parallel) or against (perpendicular) the direction of Earth’s magnetic field. Direction and magnitude were determined using Physics Toolbox Suite.

Temperature gradients were measured using an OMEGA HH506RA Multilogger Thermometer with T-type probes on multiple experimental days. The top and bottom of the Faraday cage was measured and found to be approximately 1°C warmer at the top versus the bottom, on average. To determine whether the upward movement observed in our assay was due to thermotaxis, we created a reverse gradient by placing the Faraday cage on a hot plate for the duration of the experiment, ensuring that the bottom of the assay was 1°C warmer than the top at the beginning and end of the experiment and monitored the temperature throughout. All experiments were performed at room temperature (ranging from ∼22-25°C).

#### Imaging

Fluorescent micrographs were taken using a Leica SP8 Resonant Scanning Confocal microscope at 10-40x. Worms were prepared for imaging by mounting on 2% agarose pads in 10 mM levamisole. Brightness and contrast were adjusted and overlays added using ImageJ.

#### Calculations

Estimates of force due to gravity on *C. elegans* dauers were calculated based on known and experimentally determined physical parameters. The shape of a dauer larvae is roughly cylindrical with a radius and length of 10 *μ*m and 450 *μ*m, respectively ***Kim et al. (2022***).

V = *π* 10 *μ*m^2^ x 450 *μ*m = 141.4 pL. Therefore, the estimated volume of a dauer larvae is 141.4 pL. Estimates of *C. elegans* density vary based on experimental method ***Kim et al. (2022***); ***Reina et al. (2013***); assuming the maximum density of 1.09 g/mL (based on the empirically determined density of L2-L4 larvae), the force due to gravity on a dauers was approximated.

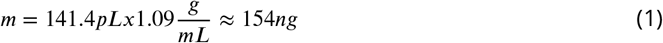

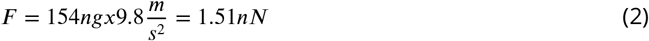

### Quantification and Statistical Analysis

Data were collected in Excel and analyzed using Rstudio. Worm positions across assays were summed and overall distributions were plotted for each strain or condition. Histograms depict the percentage of worms found at each location across all experiments. Variation between trials for Figure 2 can be found in the Supplementary Materials ***figure Supplement 2***. In some plots, the range shown along the x-axis of the histograms varies to accommodate plots in which more worms were found at one location. Boxplots provide summaries of the distributions along with statistical significance.

In addition to horizontal controls, normal distributions with a mean of 0 were simulated to compare with vertical and other assays. These distributions were created either by keeping the standard deviation (Normal Distribution^*a*^) or the number of worms (Normal Distribution^*b*^) consistent with the combined set of vertical data for each condition. Keeping the standard deviation consistent led to a slightly reduced sample size in the simulated Normal Distribution^*a*^, as the distribution is wider than the fixed range of -7 to +7; however, these distributions represent 94% or more of the total vertical sample sizes in each case. Meanwhile, Normal Distribution^*b*^ has an identical sample size to each vertical condition at the expense of narrower standard deviations. Distributions were compared using Kruskal-Wallis followed by Dunn’s test and Bonferroni’s correction for multiple comparisons. In some instances, a Gravitaxis Index (or Burrowing Index) was calculated by subtracting the number of worms in the bottom of the assay from the number of worms in the top of the assay and dividing by the total 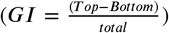. Statistical tests, p values, sample sizes, and replicates are detailed in the results and figure legends.

## Acknowledgments

Funding for this work was provided in part by NIH grants #R01GM143771 and R01HD081266 to JHR. CA was supported in part by fellowships from the Department of Molecular, Cellular, and Developmental Biology as well as the President’s Dissertation Year Fellowship through UCSB Graduate Division and an American Rescue Plan (ARP) Grant through HEERF III. LW was funded as a Dr. Rajendra Singh Fellow through the UCSB College of Creative Studies. NZK received support through the Gorman Scholarship Program at UCSB. RK received funding as a UCSB Academic Research Consortium Scholar. We acknowledge the use of the NRI-MCDB Microscopy Facility and the Resonant Scanning Confocal supported by the NSF MRI grant DBI-1625770.

Sequencing was performed by the DNA Sequencing Facility at UC Berkeley. Statistical consultation was provided by the DATALAB in the UCSB Department of Statistics and Applied Probability.

Some strains were provided by the Caenorhabditis Genetics Center, which is funded by NIH Office of Research Infrastructure Programs (P40 OD010440).

We are grateful for the generous gifts of transgenic worms provided to us by the Treinin Lab (Hebrew University), Chalfie Lab (Columbia University), and Shen Lab (Stanford University).

## Author contributions

**CA:** Conceptualization, Methodology, Formal analysis, Investigation, Writing – Original Draft, Visualization, Supervision. **LW:** Methodology, Validation, Investigation. **NZK:** Methodology, Validation, Investigation. **RK:** Investigation, Formal Analysis. **ZS:** Methodology, Validation, Investigation. **VD:** Methodology, Validation, Investigation. **KC:** Methodology, Validation, Investigation. **EL:** Methodology, Validation, Investigation. **CW:** Conceptualization, Methodology. **ES:** Methodology, Validation, Investigation. **GP:** Methodology, Investigation. **MS:** Project administration. **PJ:** Conceptualization, Writing – Review & Editing. **JR:** Conceptualization, Writing – Review & Editing, Project administration, Funding acquisition.

## Declaration of Interests

The authors declare no competing interests.

## Notes

### Competing Interest Statement

The authors have declared no competing interest.

### Summary of Updates

Additional experiments were performed to determine the effects of Earth's magnetic field on vertical preference. Additional strains with mutations in DEG/ENaC/TRN-related genes were assayed for their ability to gravitax. Several figures have been updated and/or added including six new supplemental figures as well as 10 new sub-panels to existing figures. Results and Discussion have been expanded and main text has been edited for clarity. Several authors have been added.

