## Supplemental Figures for "Mechanosensory Systems and Sensory Integration Mediate *C. elegans* Negative Gravitaxis"

### Supplemental Figure Legends

**Figure 2—figure supplement 1.** Faraday cage integrity influences gravitactic behavior

Plot of Gravitaxis Index (GI) – calculated as the sum of worms in the top half of the chamber (segments +1 to +7) minus worms in the bottom half (-1 to -7) divided by the total - for all chambers across different experimental days. Calculated for N2 dauers only. Dashed line indicates reinforcement of the Faraday cage.

**Figure 2—figure supplement 2.** Solenoid coil assay setup

**(A)** Diagram of the solenoid coil used to measure *C. elegans* response to magnetic field pertubations in the gravitaxis assay. At the University of California, Santa Barbara where these experiments were performed, the strength and direction of the magnetic field were measured using the Physics Toolbox Suite app for iOS and Android (Methods). The solenoid coil was used to counteract the vertical component of this field with measurements from a test run shown in **(B)**.

**Figure 2—figure supplement 3.** Worms oriented parallel or perpendicular to Earth's magnetic field also gravitax

**(A)** Diagram representing the orientations of gravitaxis assays for this experiment. "Parallel" tubes were oriented at 58° from the ground, which was parallel to the local measured electromagnetic field (see Methods for details). Conversly, "perpendicular" tubes were oriented 90° relative to the electromagnetic field. **(B)** Histograms depicting vertical, parallel, and perpendicular movement compared with simulated normal distributions. **(C)** Boxplots summarizing data shown in A. Parallel n = 470 worms over 6 trials; perpendicular n = 400 worms over 5 trials. $*$ $p < 0.05$, $**$ $p < 0.01$, $***$ $p < 0.001$ , $****$ $p < 0.0001$; n.s. is not significant using Kruskal-Wallis followed by Dunn's test with Bonferroni correction. Notches on boxplots represent 95% confidence intervals; mean values are indicated with a diamond.

**Figure 2—figure supplement 4.** AB1 dauers isolated in Adelaide, Australia migrate up despite inverted local magnetic field inclination

**(A)** Histograms depicting horizontal and vertical movement of N2 and AB1 dauers. **(B)** Boxplots summarizing data shown in A. AB1 vertical n = 775 worms over 8 trials. $*$ $p < 0.05$, $**$ $p < 0.01$, $***$ $p < 0.001$ , $****$ $p < 0.0001$; n.s. is not significant using Kruskal-Wallis followed by Dunn's test with Bonferroni correction. Notches on boxplots represent 95% confidence intervals; mean values are indicated with a diamond. **(C)** Insets from the US/UK World Magnetic Model – Epoch 2020.0 Main Field Inclination (I) map developed by NOAA/NCEI and CIRES (available at https://ngdc.noaa.gov/geomag/WM). Red and blue lines indicate positive and negative magnetic field inclinations, respectively. Stars have been added to show Bristol, England and Adelaide, Australia where N2 and AB1 worms were each originally isolated.

**Figure 2—figure supplement 5.** A negative gravitactic trend is also observed in *Landler et al., 2018*

Data reprinted and new statistics run with permission from the authors. In the original paper, burrowing behavior was measured in an artificial magnetic field similar to the solenoid coil experiments presented in this study. The burrowing index for each trial (equivalent to the gravitaxis index) have been plotted with circles representing fed adult N2 worms, triangles representing starved worms, and blue and grey symbols representing worms exposed to positive and negative field inclinations, respectively. A two-sided t-test was performed by the authors of this study and annotated accordingly (∗ p <0.05).

**Figure 2—figure supplement 6.** Gravitaxis behavior occurs independently from thermotaxis along shallow temperature gradients

**(A)** Histograms depicting horizontal and vertical movement of N2 dauers along normal and reverse temperature gradients. **(B)** Boxplots summarizing data shown in **A**. Reverse gradient n = 445 worms over 5 trials. ∗ p <0.05, ∗∗ p <0.01, ∗∗∗ p <0.001 , ∗∗∗∗ p <0.0001; n.s. is not significant using Kruskal-Wallis followed by Dunn's test with Bonferroni correction. Notches on boxplots represent 95% confidence intervals; mean values are indicated with a diamond.

**Figure 4—figure supplement 1.** Gravitaxis behavior in additional strains

**(A)** Histograms depicting horizontal and vertical movement of N2 and LSJ1 worms as well as additional mechanosensory deficient dauers. *mec-4(u253)* and *mec-6(e1342)* dauers were found to be sluggish, resulting in little taxis in either direction. †*mec-6* dauers accumulated at the origin of the assay; the x-axis was altered in plot to allow for visual comparison. Additional alleles of *mec-7* and *mec-10* recapitulate the findings shown in results. **(B)** Boxplots summarizing data shown in **A**.LSJ1 vertical n = 1,666 worms over 7 trials; horizontal n = 2,510 worms over 7 trials. mec-4(u253) vertical n = 2,075 worms over 7 trials; horizontal n = 2,130 worms over 4 trials. mec-6(e1342) vertical n = 195 worms over 6 trials; horizontal n = 277 worms over 4 trials. mec-7(e1343) vertical = 949 worms over 8 trials. mec-10(e1515) vertical n = 1,244 worms over 4 trials; horizontal n = 1,334 worms over 2 trials. ∗ p <0.05, ∗∗ p <0.01, ∗∗∗ p <0.001 , ∗∗∗∗ p <0.0001; n.s. is not significant using Kruskal-Wallis followed by Dunn's test with Bonferroni correction. Notches on boxplots represent 95% confidence intervals; mean values are indicated with a diamond.

**Figure 4—figure supplement 2.** Worm distribution variation between trials

Histograms depicting vertical movement of N2 worms as shown in Figure 2. SEM is plotted at each vertical location to show variation in worm location between trials. Sample sizes and trial numbers are listed in the table below.

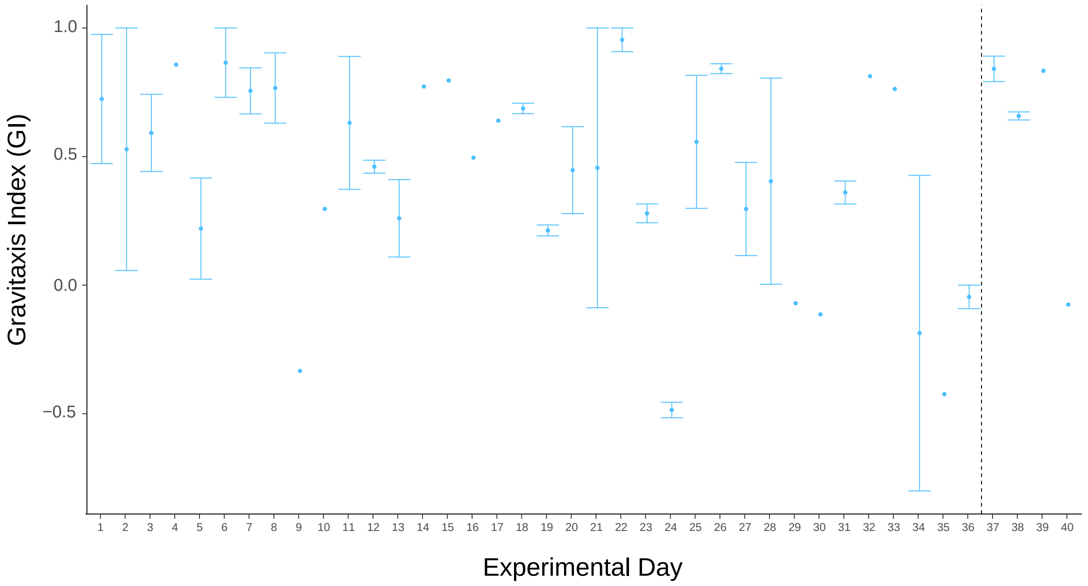


**Figure 2—figure supplement 1.** Faraday cage integrity influences gravitactic behavior

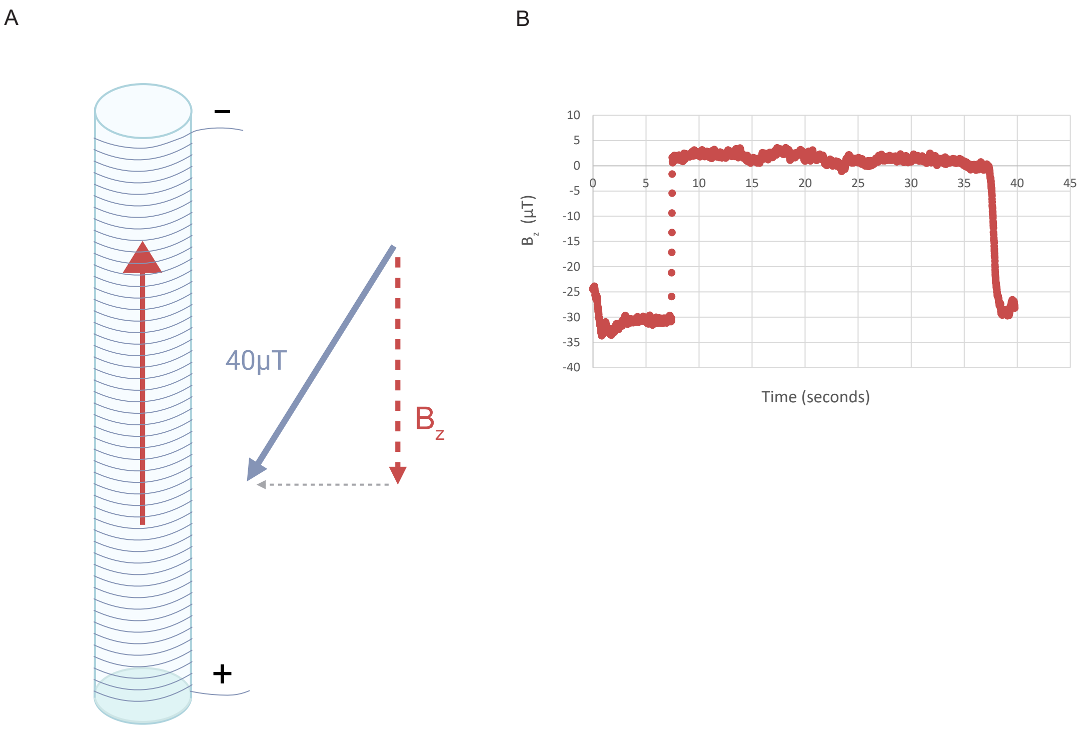


**Figure 2—figure supplement 2.** Solenoid coil assay setup

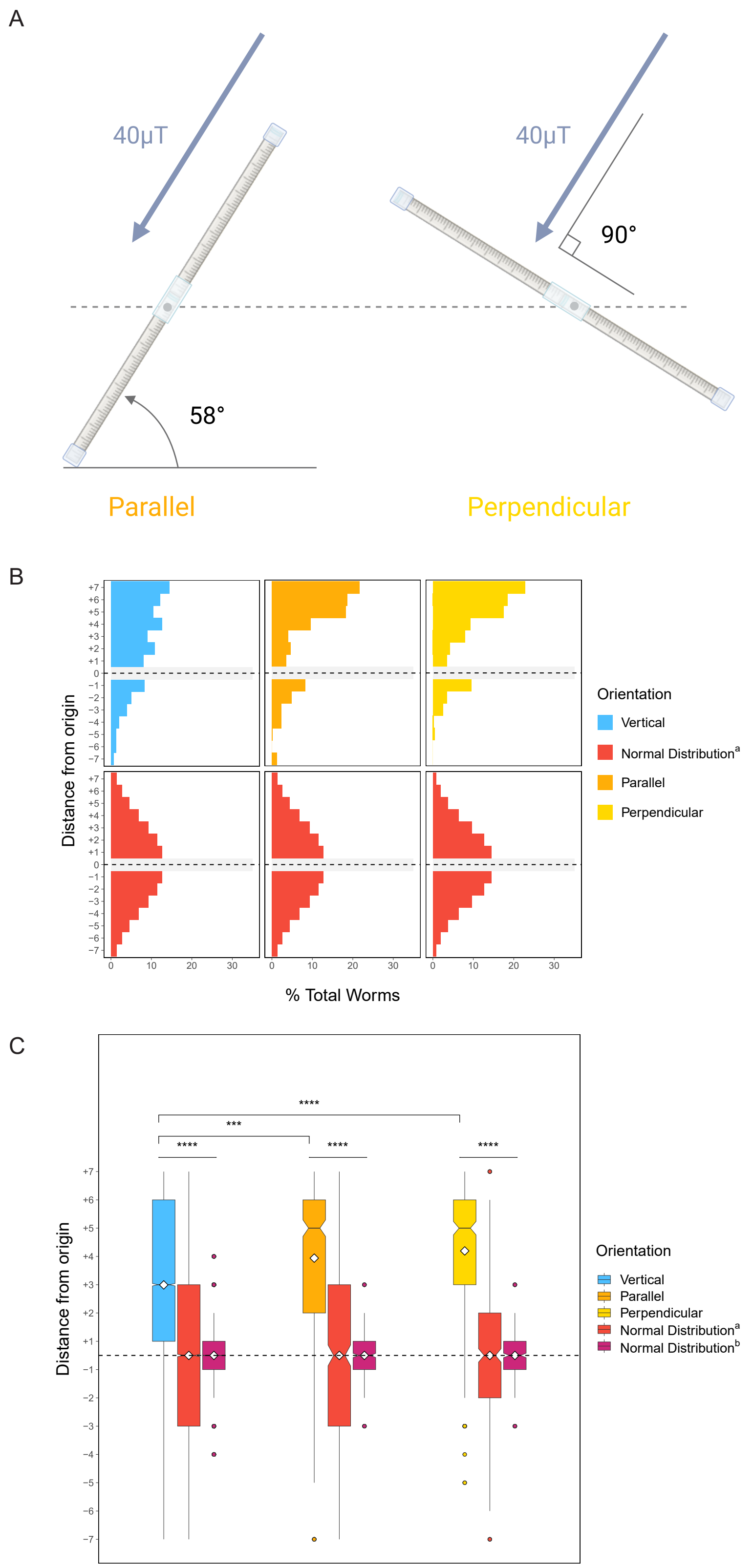


**Figure 2—figure supplement 3.** Worms oriented parallel or perpendicular to Earth's magentic field also gravitax

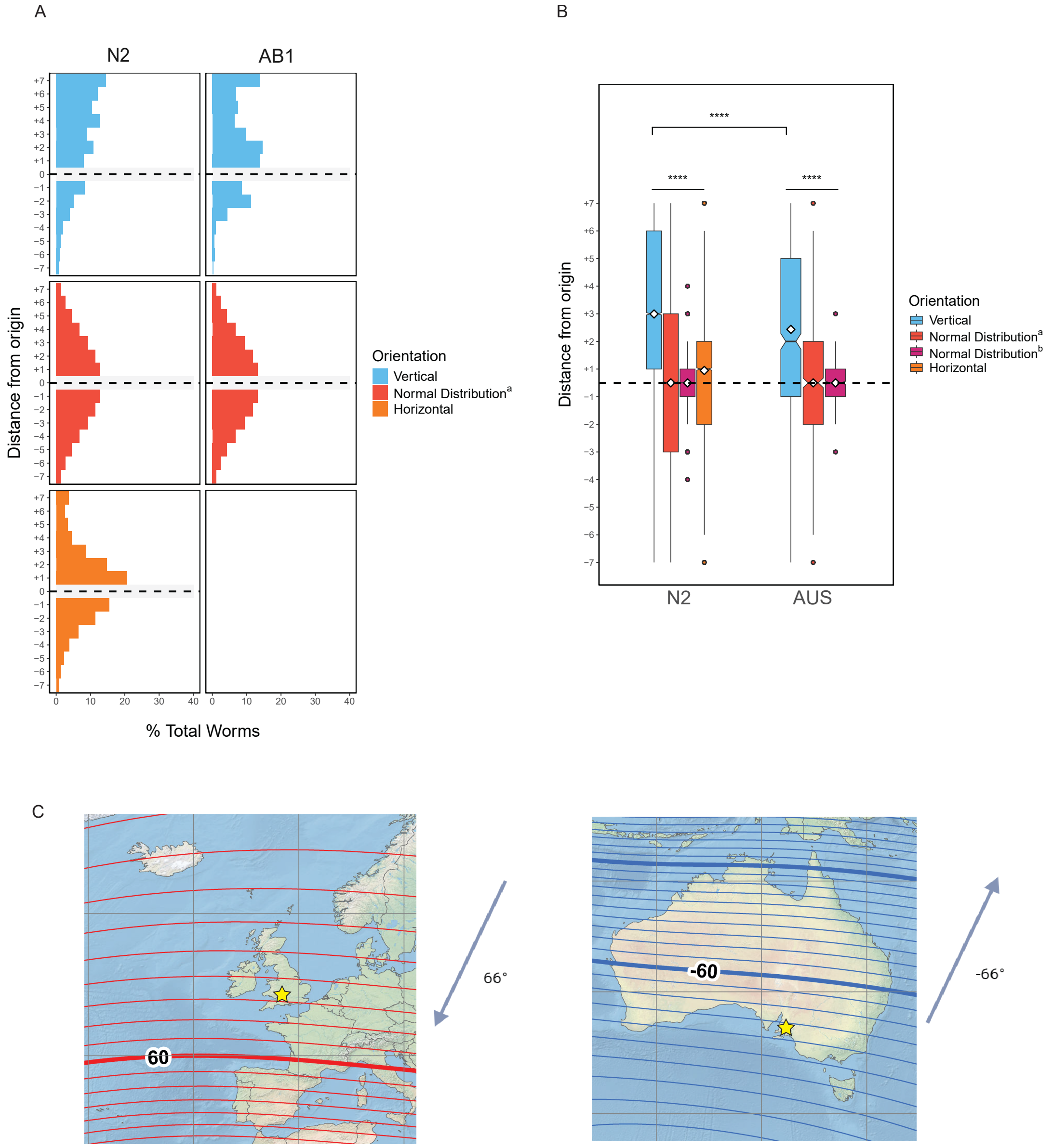


**Figure 2—figure supplement 4.** AB1 dauers isolated in Adelaide, Australia migrate up despite inverted local magnetic field inclination

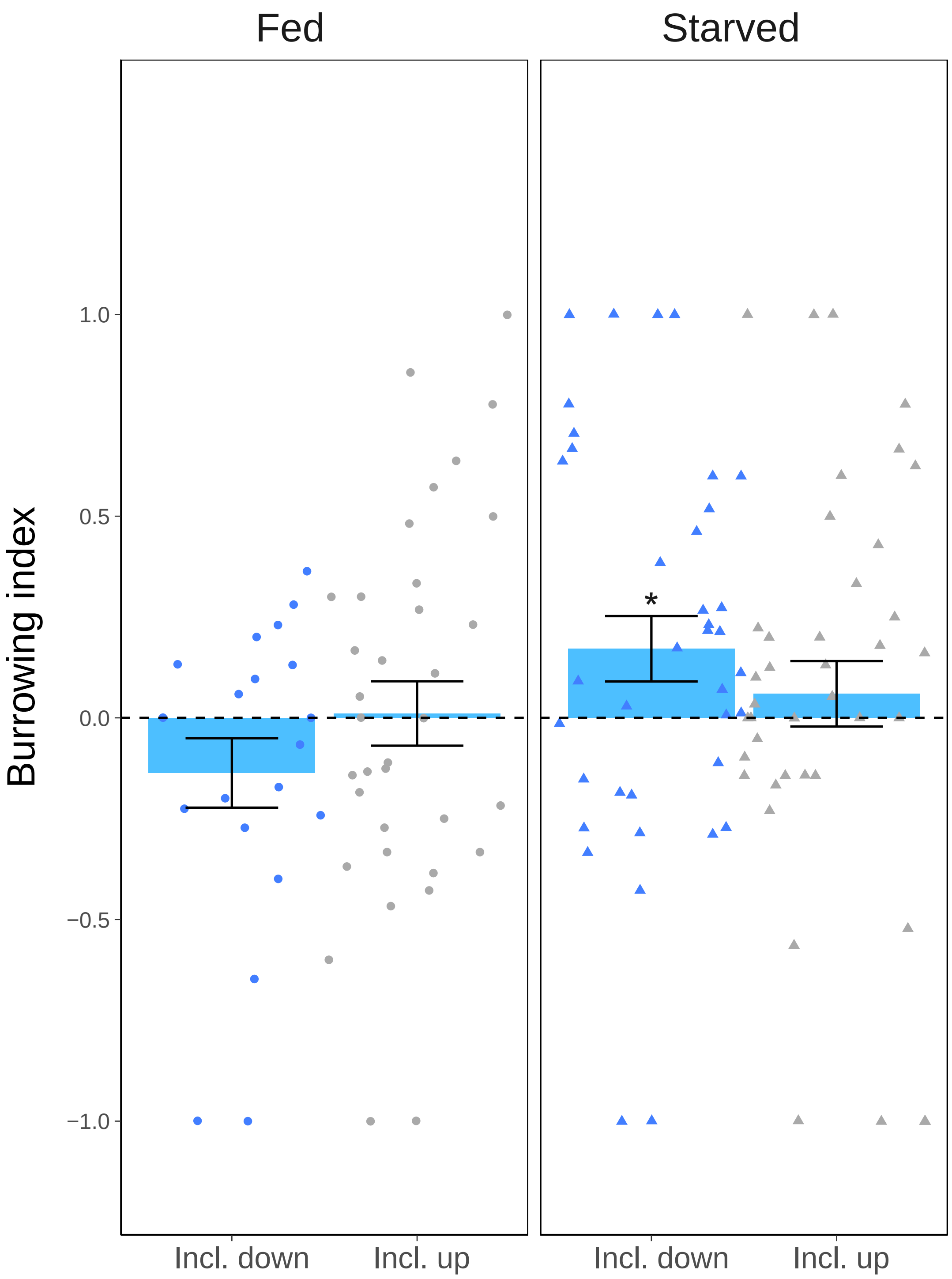


**Figure 2—figure supplement 5.** A negative gravitactic trend is also observed in ***Landler et al. (2018b)***

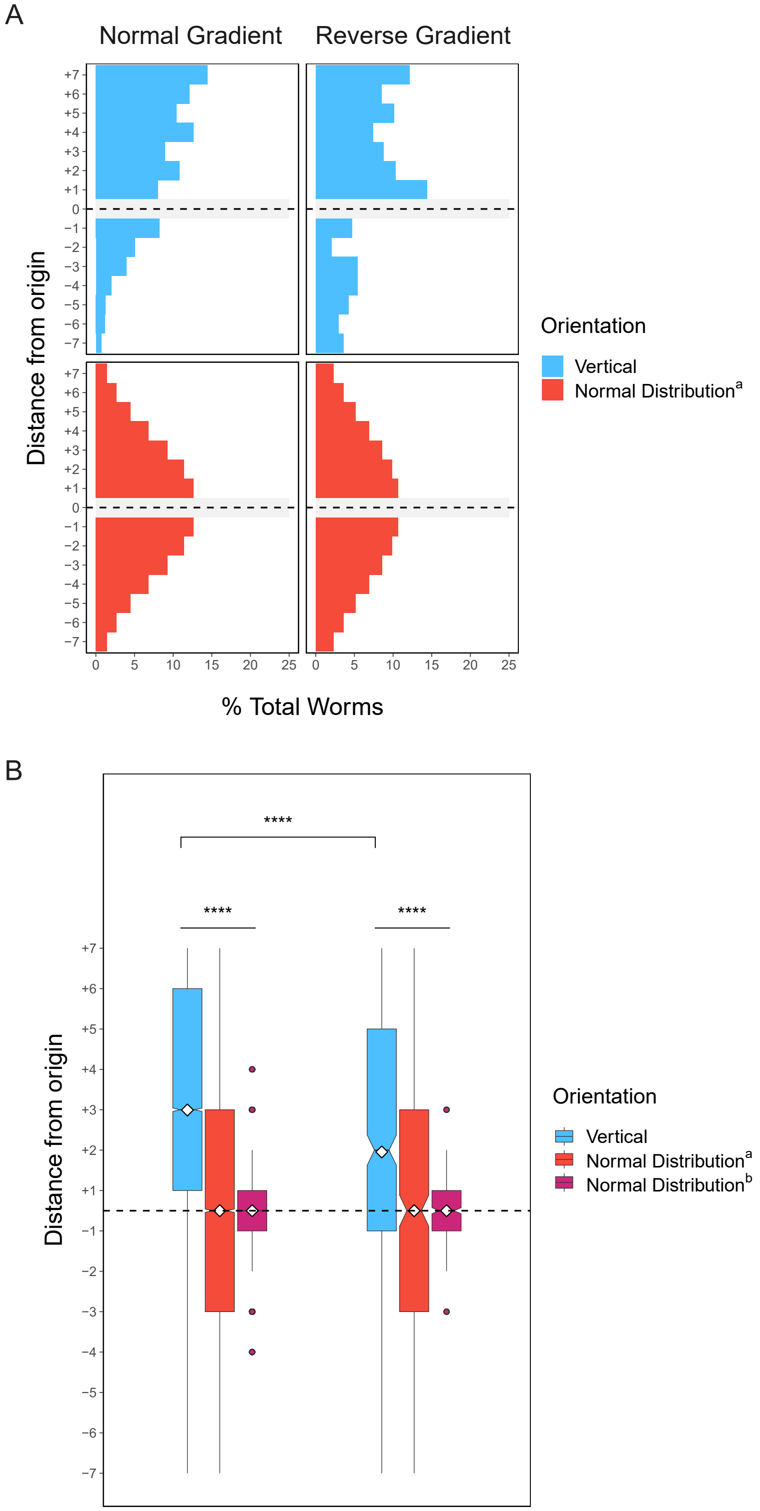


**Figure 2—figure supplement 6.** Gravitaxis behavior occurs independently from thermotaxis along shallow temperature gradients

1107

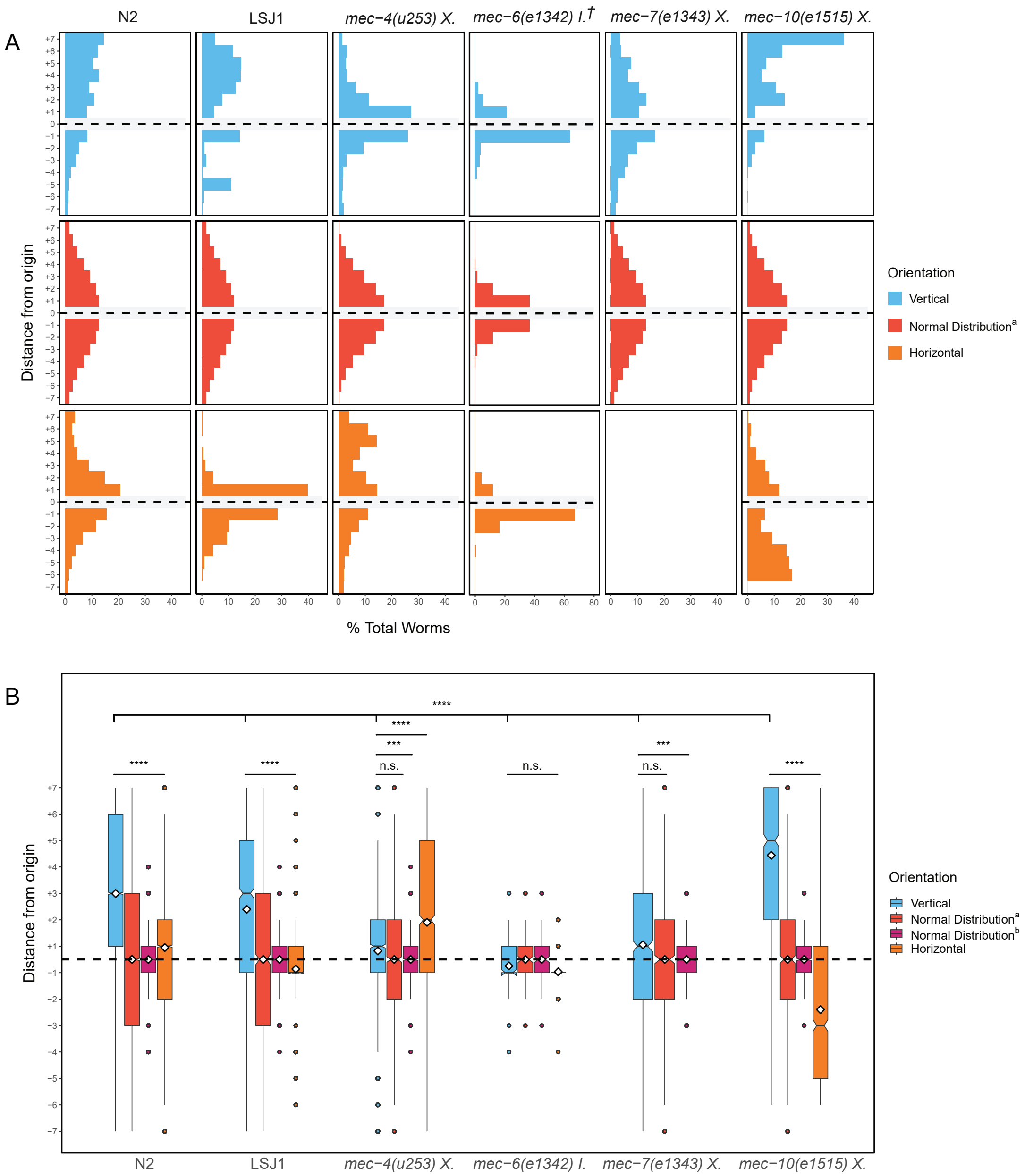


**Figure 4—figure supplement 1.** Gravitaxis behavior in additional strains

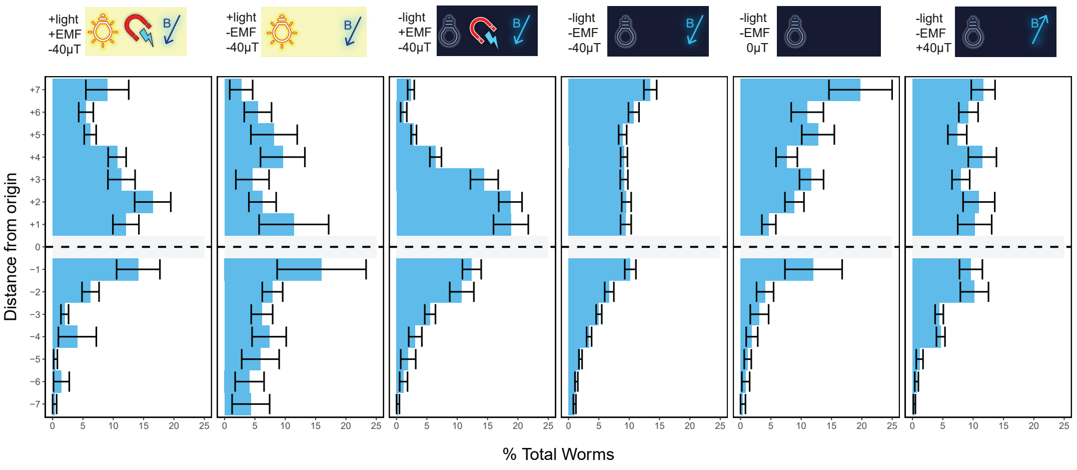


| | | | | | | |
|---|---|---|---|---|---|---|
| n | 2222 | 542 | 4358 | 35816 | 696 | 1112 |
| trials | 10 | 6 | 12 | 148 | 9 | 9 |

**Figure 4—figure supplement 2.** Worm distribution variation between trials
